# Sex differences in development alter the fledgling sex ratio in a lekking bird with strong sexual size dimorphism

**DOI:** 10.1101/2025.10.29.685087

**Authors:** Lina M. Giraldo-Deck, James D. M. Tolliver, Hanna Algora, Jelena Belojević, Luke J. Eberhart-Hertel, Bart Kempenaers, Kari Koivula, Krisztina Kupán, Katrin Martin, Veli-Matti Pakanen, Veronika A. Rohr-Bender, Nelli Rönka, David B. Lank, Clemens Küpper

## Abstract

Sex ratio variation is of fundamental importance for population ecology, and the evolution of sex roles and life-histories. Yet, the ecological mechanisms underlying such variation remain often unknown. Using a multiyear dataset from captive and wild precocial Ruffs *Calidris pugnax*, we show that poorer survival and later fledging of males relative to females result in a female biased sex ratio of the fledgling population. Both sex-specific survival and maturation time contributed equally to an increasing sex ratio bias from hatching to fledging in this sexually dimorphic bird. On average, juvenile females fledged three days earlier than males due to expedited wing development, despite the higher absolute rate of wing growth in males. We argue that sex differences in growth, maturation, and offspring survival are likely the result of strong pre-sexual selection on males to achieve dominance and may help to enforce the Darwinian sex roles observed in this species.

## Introduction

The sex ratio (SR) is a key demographic parameter for evolutionary and ecological processes within populations. A biased SR at maturation can have important implications for population dynamics (Le Galliard *et al*. 2005; Eberhart-Phillips *et al*. 2017; Heinsohn *et al*. 2019; Chiba *et al*. 2023), sexual selection and sex roles (Kokko & Jennions 2008; Liker *et al*. 2013; Kappeler *et al*. 2023), life histories and social behaviours (Boomsma & Grafen 1990; Kappeler 2017). The “Fisher Condition” predicts a 1:1 male-female primary sex ratio when producing either sex is equally costly for the parents, because frequency-dependent benefits favour parents that produce the rarer sex (Düsing 1884; Fisher 1930; Kokko & Jennions 2008; Jennions & Fromhage 2017). However, whether parents can adaptively adjust the sex of their offspring, or the observed deviations are due to stochasticity, is debated (Trivers & Willard 1973; Krackow 2002; West & Sheldon 2002; Schindler *et al*. 2015; Eberhart-Phillips *et al*. 2017), and the mechanisms shaping SR variation remain poorly understood.

The primary SR, measured at conception, is typically unbiased, implying that a biased adult SR will develop post-conception (Székely *et al*. 2014; Jennions & Fromhage 2017). Biasing factors include differential mortality for males and females either before or after they reach the adult stage (Eberhart-Phillips *et al*. 2017; Regan *et al*. 2020; Eberhart-Hertel *et al*. 2024), different maturation times with the earlier maturing sex becoming more frequent in the adult population (Lovich & Gibbons 1990; Hirst *et al*. 2010; Staufer *et al*. 2023), and sex-specific dispersal. Variation in mortality, maturation, and movement often relates to divergent sex roles during adult life. For example, providing care may increase parental mortality risk, resulting in the sex that engages more in care becoming scarcer (Jennions & Fromhage 2017). Yet, this mortality bias can be reversed if members of the less-caring sex invest more in sexually selected traits that increase mortality costs to their bearers. Such traits can be expressed temporarily or continuously. Temporarily expressed traits include ornaments, bright colouration, or elaborate songs that make their bearers more attractive to the opposite sex during the mating season but also make them more vulnerable to predators, hence increasing the mortality risk (Andersson 1994). Continuously expressed traits include body size, where large size often confers advantages for fertility or higher competitive ability in dominance contests but requires substantial resources and time for growth and maturation. Consequently, large individuals have higher growth rates during development and/or take longer to mature than smaller individuals (Ricklefs 1979; Richner 1991; Blanckenhorn *et al*. 2007; Nijhout *et al*. 2010).

For species with determinate growth, size differences between sexes and among individuals develop early during ontogeny. Any costs associated with large size will start to accrue during this time (Badyaev 2002) and may result in a mortality bias that widens with increasing sexual size dimorphism (Clutton-Brock *et al*. 1985; Promislow 1992; Benito & González□Solís 2007). Since males are often the larger sex, higher pre-adult mortality in males is common, but a positive size-dependent mortality relationship is also observed in species where females are larger than males (Torres & Drummond 1997; Kalmbach *et al*. 2005; Loonstra *et al*. 2019). Environmental conditions may affect the survival of the two sexes differently, with the larger sex being more sensitive to resource shortage than the smaller sex because large individuals require more energy to reach their final size (Anderson *et al*. 1993; Kalmbach *et al*. 2005; Loonstra *et al*. 2019).

Differences in development time may also contribute to sex-specific mortality. Prolonged development will increase mortality of juveniles when individuals are more vulnerable to predation at earlier rather than at later life-stages. This is particularly the case when locomotive ability increases with development. In many birds, obtaining flight ability (i.e., “fledging”) represents an important milestone for juveniles on their way to independence. Fledging enhances the locomotive ability and can improve survival as it allows juveniles to escape ground predators more efficiently (Schekkerman *et al*. 2009). Yet, there is considerable variation in fledging ages among and within bird species (Martin *et al*. 2018). Sex-specific life-history requirements may contribute to this variation but so far remain unexplored. Sex differences in fledging age could signal a classic life-history trade-off where early fledging increases survival whereas late fledging increases future reproductive success. This trade-off arises because a larger body size, which is often favoured in male-male contest competition or for female fertility, typically requires longer development time (Badyaev 2002). In addition, a larger mass requires higher force to lift the bird off the ground and hence larger and better developed wings (Alerstam *et al*. 2007). Similar to the observed variation in fledging ages, there is considerable variation in wing development between and within species (Jones *et al*. 2020a). However, it remains unclear to what extent within-species variation in wing development and maturation is sex dependent.

The Ruff *Calidris pugnax*, is a migratory lekking sandpiper that breeds in wetlands and the tundra of the northern Palearctic. Because of (1) the large sexual dimorphism fuelled by intense male-male competition (van Rhijn 1991), and (2) a biased SR at the time of fledging (Jaatinen *et al*. 2010), Ruffs provide a highly appropriate study species to test how SR biases develop during early ontogeny. Adult males are on average ∼70% heavier than females (Lank *et al*. 2013; Giraldo-Deck *et al*. 2020). At hatching, the precocial male and female chicks are the same size. However, males grow faster than females resulting in a clear size dimorphism already detectable about one week after hatching (Giraldo-Deck *et al*. 2020). The size differences between adult males and females are tied to Darwinian sex roles where males compete over dominance on the lek, whereas females are choosy and provide substantial resources, including all parental care (Hogan-Warburg 1966; van Rhijn 1991; Giraldo-Deck *et al*. 2022). A strong mating skew among males has led to the evolution of three specialised reproductive morphs – Independents, Satellites and Faeders (Hogan-Warburg 1966; Jukema & Piersma 2006; Lank *et al*. 2013) – that are determined by an autosomal inversion polymorphism that arose around 4 MYA (Küpper *et al*. 2016; Lamichhaney *et al*. 2016). Ancestral Independents are the most common morph, representing about 80-90% of the adult population (Hugie & Lank 1997; Widemo 1998; Lamichhaney *et al*. 2016). Independents are characterised by strong and aggressive dominance contests among males, which likely promoted the evolution of strong sexual size dimorphism (Darwin 1871; Oakes 1992; Parker 1992). In contrast, the reproductive tactics of the other two morphs, Satellites and Faeders, which are on average smaller than Independents (Lank *et al*. 2013), are not based on competition over dominance.

Here, we examine how pre-fledging development and mortality contribute to SR alteration from hatching to fledging. We focus exclusively on Independents, as they are the most common and ancestral morph. First, using a large dataset on individual survival and age-at-fledging from Ruffs that were hand-raised in captivity, we compare fledging ages between male and female juveniles and determine how fledging age relates to body size. Second, we investigate variation in wing growth and feather maturation between males and females as a potential mechanism for the observed differences in fledging age. Third, we use a population matrix model parameterized with fledging and survival data from the aviary, as well as empirical data on the hatching SR, and on predation and survival of individually marked chicks in the wild to characterize the phenology of SR changes until fledging. Taken together, our results reveal how sex differences in growth, maturation time, and survival generate a burgeoning SR bias before maturation that is likely driven by intense competition for dominance in adult males.

## Material and Methods

### Development and fledging of captive

*Ruffs.* From 2019 until 2023, we monitored wing growth, feather maturation and flight ability of captive Ruffs of the Independent morph (N=196 fledglings from 88 mothers). Measurements of body mass, wing length, tarsus length and feather growth (i.e. the length of the longest primary) were taken at two locations, Burnaby (Canada) and Seewiesen (Germany; see Supplementary Methods). The population was originally established in Canada from eggs collected near Oulu, Finland (Lank *et al*. 2013). It was translocated to Seewiesen in 2019 and 2020 and supplemented with additional Ruffs obtained from breeders and zoological gardens in the Netherlands, Belgium and Germany.

To assess flight ability, we conducted daily flight tests starting when a bird became 16 days old. For the test, the prospective fledgling was placed on a box (height: approx. 50cm), which was standing in front of a water bath (approx. 2.0m x 2.0m). A person standing behind the box prevented the test bird from jumping off the box at either side and gently nudged it towards the water. Birds that flew over the water bath passed the test, whereas birds that hopped into the water failed it. To determine the efficacy of the test, we continued flight tests in a subset of individuals (N=15) for two further days after their first successful flight: all 15 individuals passed their subsequent trials and flew over the water bath again. Consequently, we defined fledging age as the age at which an individual first passed the test.

To examine differences between the sexes in fledging age and the effect of adult body size, on fledging age, we compared posterior means and 95% credible intervals (CrIs) from two linear models with fledging age as response variable. As measures for adult body size, we used body mass or tarsus length measured in the first or second winter. The first model contained sex and adult body mass as fixed effects, and cohort (hatch year) and mother identity as random effects. The second model had a similar structure, but instead of adult body mass, we used adult tarsus length as a fixed effect.

To compare wing length, wing growth and feather maturation in juveniles between sexes we first modelled posterior means of wing or emerged feather length (and their 95% credible intervals (CrIs) in relation to age for each sex using the measurements that were taken two to three times per week (see Supplementary Material). For these models, we used the R package BAMLSS, a flexible Bayesian generalized additive model framework (GAMM) based on Markov chain Monte Carlo simulations that can include random factors (Umlauf *et al*. 2018). We modelled each sex separately because categorical fixed effects cannot be specified in BAMLSS and our main interest was to examine differences in the shape of growth curves between males and females. For each model, we z-transformed the variables wing length, feather emergence and age by setting their means to zero and their standard deviation to one. We included ‘mother ID’, ‘Individual ID’ and an interaction between ‘Individual ID’ and ‘age’ as a random factor in all models to allow for individual specific growth curves. In separate models, we estimated posterior means and 95% CrI of wing growth rates, the proportion of adult wing length, the rate, and proportion of feather emergence in relation to age.

For each model, we obtained model estimates through simulation of 10,000 values from the joint posterior distribution of the model parameters using the function ‘sim’ of the package arm (Gelman & Su 2015) and a flat prior distribution. We report model estimates (means) with their 95% CrI (the 2.5% and 97.5% quantiles of the simulated values) and provide posterior probabilities for specific hypotheses. Probabilities higher than 0.975 or lower than 0.025 would indicate statistical significance at an α-threshold of 0.05 according to more traditional frequentist statistics. To assess model fit, we analyzed residual distributions graphically using normal quantile-quantile plots and by plotting residuals against leverage and against fitted values of each factor. For all analysis, we used R version 4.4.2 (R Development Core Team 2024).

### Data sources for Population Matrix Model

To investigate changes in SR from hatching to fledging, we constructed a population matrix model that we informed with three distinct datasets. First, hatching sex ratio *ρ* and chick survival data, collected from 313 clutches (Tables S1, Fig. S2) and from 36 radio-tagged broods of free-living Ruffs (Fig. S3). As others (Eberhart-Phillips *et al*. 2018), we used *ρ* as a proxy of the primary sex ratio, and assumed no sex difference in embryonic survival. Second, sex- and age-specific survival and fledging data, obtained from 377 and 196 captive Ruffs, respectively (Table S2, Fig. S4, S5). Third, published age-specific survival estimates of wader chicks before and after fledging (Schekkerman *et al*. 2009) – as comparable data for Ruffs are not available – to derive the ‘fledgling advantage’ *F* (Fig. S6). Predation is a major source of mortality in wader chicks (Mason *et al*. 2018). Because captive chicks were raised safe from predators, we emulated a more natural scenario and implemented an ‘additive predation mortality’, defined as the observed age-specific survival differences between captive and wild (radio-tracked) chicks (Supplementary Methods).

To quantify *ρ* and estimate chick survival in the wild, we collected data from a breeding population at Pitkänokka meadows, Liminka Bay, Finland (64°85‘93.0"N, 25°27’61.0"E) from 2016 until 2023. Information on nest searching and monitoring is provided elsewhere (Algora *et al*. 2025; Koivula *et al*. 2025). Soon after hatching, we weighed chicks, collected a small blood sample (20–75 µl) from the tarsal vein for molecular sexing (Giraldo-Deck *et al*. 2020), and marked each chick with a numbered metal ring. We modelled the *ρ* with a binomial mixed effects model where our response variable was the probability of a hatched chick being a male (0, 1). We first fitted z-transformed relative clutch initiation date (‘Season’) as a fixed effect, and ‘Clutch ID’ and sample year (‘Cohort ID’) as random effects (Table S3). Because ‘Season’ had no clear effect on the HSR, we omitted it from the final model (Table S3).

### Matrix model structure

To examine how the effects of survival and fledging age alter the SR from hatching through fledging, we constructed a two-sex population matrix model with sex-specific parameters (Fig. S1). We used a Bayesian statistical framework for estimating model parameters and informed the matrix model with the posterior distributions of the parameter estimates and projected the population through an array of sex and age-specific matrices, where each matrix represented a single day in the life of a chick (Supplementary Methods).

### Predicting post-fledging SR

We estimated the post-fledge SR by projecting cohorts of Ruff chicks from hatching (*t*=0 days) to the age when all chicks had fledged (*t*=27 days). We started simulations with 100,000 hatchlings. To propagate error from our empirical estimates in vital rates, we simulated cohorts in an analogous way as we estimated posterior distributions: we generated four chains of 3,500 Ruff cohorts with 1,000 warm up simulations, resulting in a posterior distribution derived from 10,000 simulations. Each cohort had a unique combination of random draws from each of the posterior distributions of *ρ*, intrinsic survival *ϕ′* additive predation mortality *D*, and fledging probability *ψ*. We truncated the posterior distribution of *D* at 0.2 because higher values led to extinction of the simulated cohorts. The resulting posterior distribution of *D* had a mean of 0.08 (95% CrI: 0.00–0.18). We based all simulations on a fledging advantage *F* of 0.05, i.e. a 5% increase in survival after fledging (Supplementary Methods, Fig. S6).

For each of the 10,000 simulations, we calculated the post-fledge SR as *n*/(*n* + *n*), where *n* is the number of males and *n* the number of females within the post-fledge cohort at *t=27* days, the first age when all juveniles had passed the fledging tests. We calculated the difference in SR *ΔSR* between hatch and post-fledge cohorts, as well as the relative fitness and selection coefficients for each simulation. Sex-specific relative fitness *w* is the ratio of the sex frequencies at post-fledge and hatch. The cross-sectional selection coefficient *s* was calculated as Δ*p*/*p*_post-fledge_(1–*p*_hatch_), where *p* is the population proportion of the sex of interest (Linnen & Hoekstra 2009; Svensson *et al*. 2019). We report mean values and 95% CrIs across all simulations.

### Life Table Response Experiments

We used two life table response experiments (LTREs, Caswell, 2001) to examine how perturbations of our model parameters influence our model responses (post-fledge SR, *w*, and *s*). First, to quantify the relative impacts of sex-specific differences in fledging age *ΔAge* and *ϕ′* we used an alternative demographic model (‘unisex model’) to the model presented above (‘two-sex model’). For the unisex model, we kept intrinsic survival *ϕ*(*t*), i.e. the posterior weighted average of intrinsic survival at age *t,* constant for all chicks regardless of their sex, whereas all other input parameters were varied in the same way as in the two-sex model. We compared the response variables (mean and variance of SR, *w*, and *s*) of the unisex model with those of the two-sex model by dividing the values from the unisex model with their respective counterparts from the two-sex model to quantify how much of each output parameter was explained by the alternative model.

Second, we used a perturbation LTRE to measure the effect of each input parameter on the post-fledge SR and the male selection coefficient *s*_♂_. Perturbation analysis allows for the quantification of the relative impact of the model’s input parameters on population-level response(s) (Caswell 2001; Eberhart-Phillips *et al*. 2017). LTREs can thus be used to decompose the relative contributions of ecological and demographic parameters, e.g. the effects of predation or sex biases in *ϕ* on the response (Veran & Beissinger 2009; Eberhart-Phillips *et al*. 2017). Within a perturbation, the difference in the response parameter between two or more ‘treatments’ is decomposed by weighting the difference by the parameter’s contribution to a response variable, its so-called ‘sensitivity’ (Supplementary Methods), and then summing sensitivities across all parameters (Caswell 2001; Veran & Beissinger 2009; Eberhart-Phillips *et al*. 2017).

We then calculated the influence of each ecological and demographic parameter on response variables in two ways. First, we calculated the relative contributions *C* of each demographic (*ρ*, cumulative intrinsic survival *ϕ′* and *ΔAge)* and ecological (*D* and *F*) input parameters to the mean post-fledge SR and *s*_♂_ (Supplementary Methods). Second, we calculated the theoretical unit change in the responses given an increase of one standard deviation of the posterior distributions of each input parameter, or, in the case of *F*, one standard deviation of the extracted dataset. This allowed us to measure the importance of each input parameter in shaping the response variables, while also accounting for the sensitivity of the input parameter on the response variable.

## Results

### Sex differences in fledging age

The age at which juvenile Ruffs passed the flight test, ranged from 16 to 27 days. Females fledged earlier than males (Fig. 1a, b; P>0.999), regardless of whether we used body mass or tarsus length as the measure of adult body size (Tables S7 and S8). The mean difference in fledging age between the sexes was 3.1 days when fledging age was modeled using the sex-specific mean adult mass or tarsus length (Fig. 1a and S7). The mean difference increased to 5.0 (3.7–6.4) days when we used the overall mean adult body mass (140 g) in the model (Table S1), and to 4.2 (2.9–5.4) days when we used the overall mean adult tarsus length (47.91 mm) (Table S7). Within each sex, heavier individuals fledged earlier than lighter individuals (Fig. 1b, P>0.99).

**Fig. 1.**
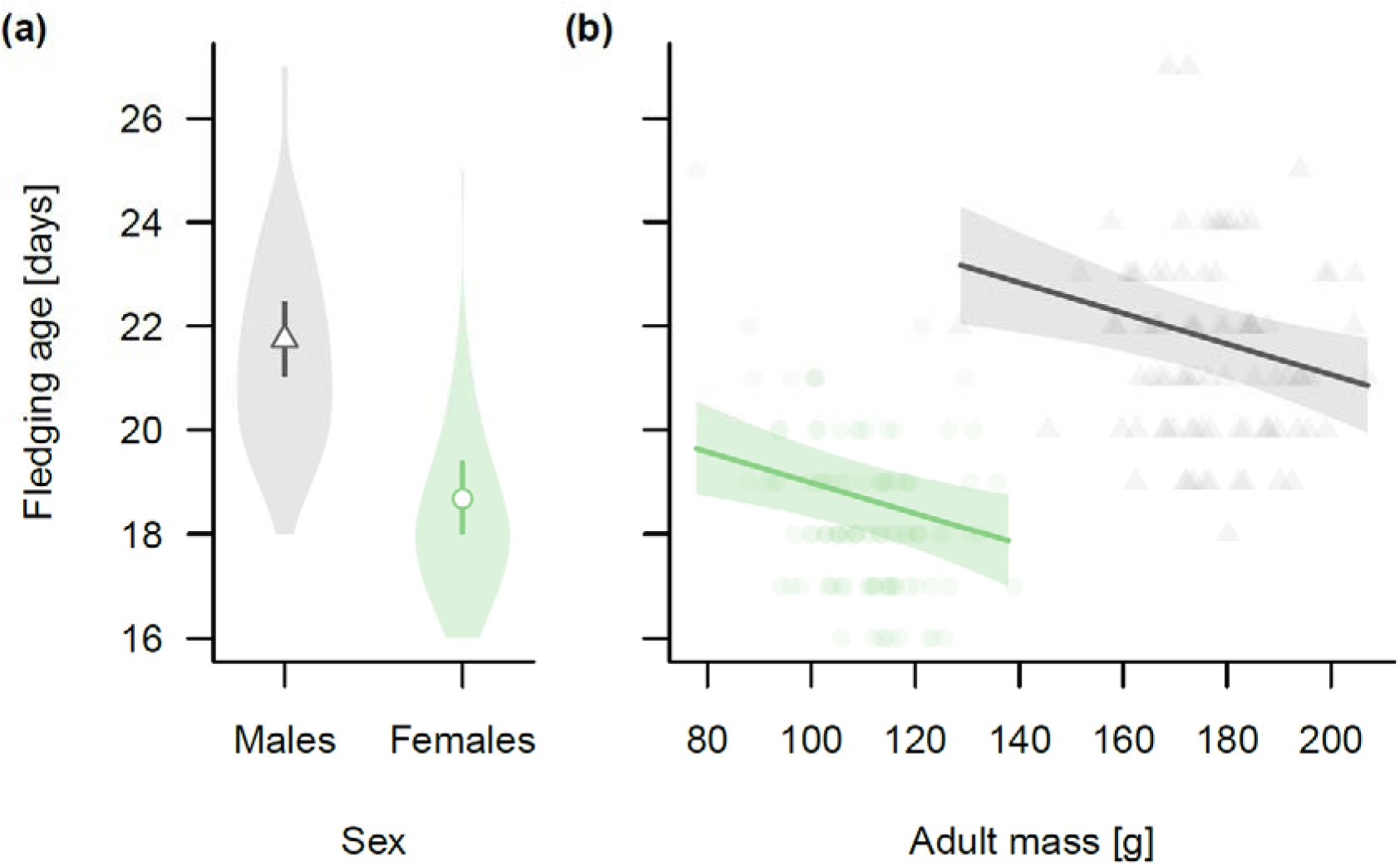
Variation in fledging age in relation to sex and adult body mass. **(a)** Sex-specific fledging ages when statistically controlling for cohort and mother ID. Females (green) fledged earlier than males (gray) (P>0.999). Means (triangle for males and circle for females) and 95% credible intervals (lines) correspond to fledging ages at a sex-specific mean adult mass (females: 110.4 g, males: 177.0 g). Shaded areas indicate the density distributions of the raw data. **(b)** Fledging age in relation to variation in adult body mass for males and females. Shown are the predicted means (lines) and 95% credible intervals (shaded area). Triangles and circles show the individual data points for males and females, respectively. Heavier individuals fledged earlier than lighter individuals within each sex (P >0.99). For full model details see Table S7.

### Sex differences in wing development

Consistent with their larger body size, the final wing length of males was larger than that of females (Fig. 2a, Table S9). Also, the pattern of growth of the wings and primaries differed between the sexes. For wing length, females reached their peak growth rate earlier than males (Fig. S8a). From day 8 onwards, the wings of males grew faster than those of females and the wing growth period was longer for males than for females, resulting in longer overall wings (Fig. 2a, Fig. S8a). The emergence of the primary feathers begun at an age of 5 days. The emergence rate and length of the primaries was initially higher for females than for males (Fig. S8b) indicating that their wings matured faster, which presumably enabled them to fledge earlier than males. At fledging, the wings of both males and females had a similar level of maturity: their wing length had reached a proportion of 0.73 of their future mature adult wings, and a similar proportion of the longest primaries had emerged (females: 0.73, males: 0.74, Fig. 2b, c, Table S9).

**Fig. 2.**
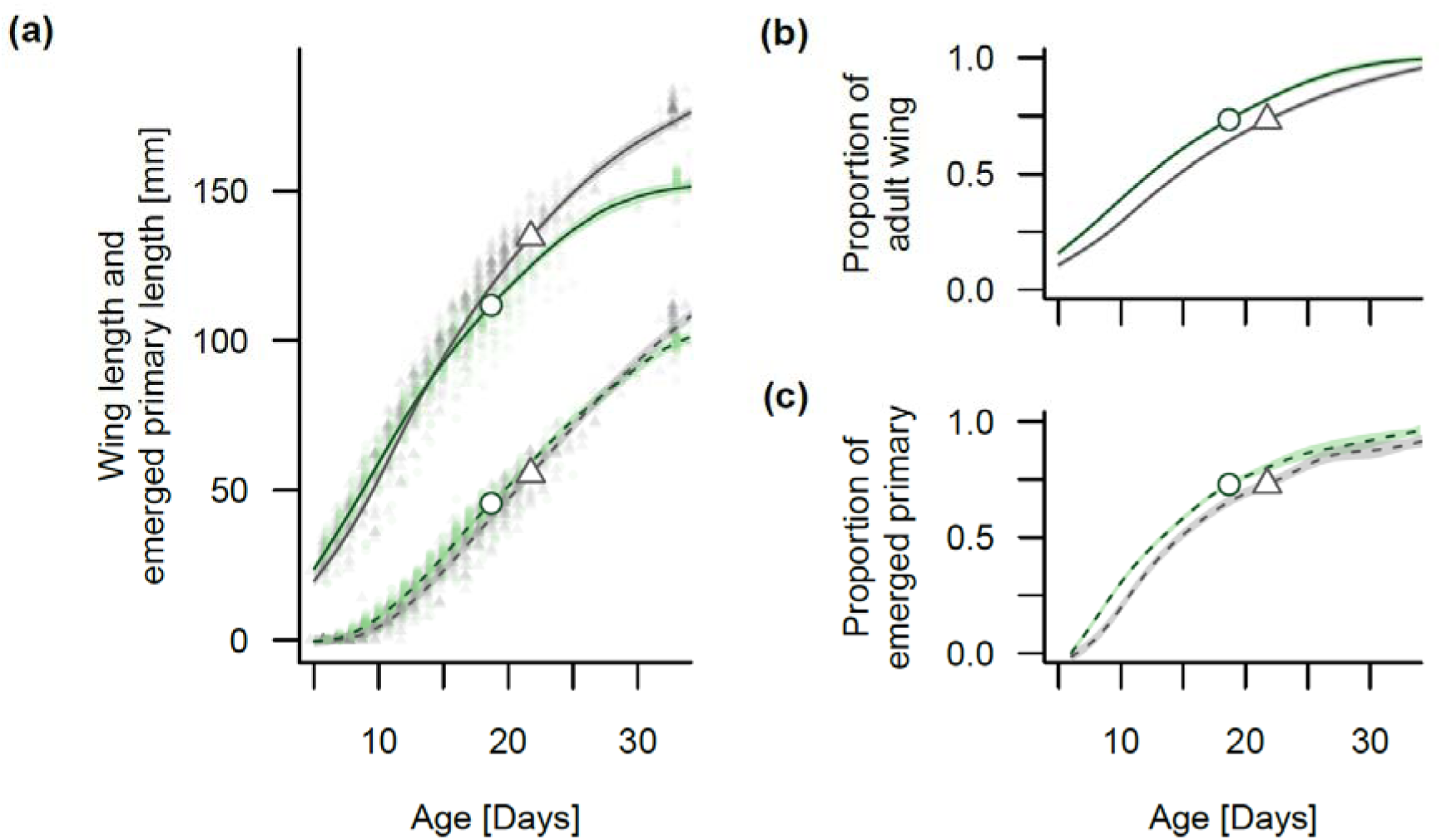
Wing development and feather maturation in males and females. (a) Growth of wing length (continuous lines) and longest emerged primary length (broken lines) in Ruff juveniles. (b) Proportion of adult wing length at a given age. (c) Proportion of emerged primary length at a given age. Shown are means (lines) and 95% credible intervals (shaded areas) of males (gray) and of females (green). Open symbols indicate mean values at fledging for males (triangles) and females (circles). Raw data are shown in (a) as light-coloured triangles or dots.

### Projection of sex ratio from hatching to post-fledging

The single-cohort model, projected a clear mean SR decline from hatching until all Ruff juveniles had fledged from 0.50 (0.46– 0.55; 95% CrI) to 0.43 (0.37–0.49; 95% CrI; Fig. 3a, P>0.999). This amounts to a 14% decline in the proportion of males in the Ruff juvenile cohort across the 27-day period. In comparison with females, males had lower cross-sectional relative fitness (mean *w*_♂_=0.86; mean *w*_♀_=1.14; Table S10) and consistently negative selection coefficients across simulations (mean *s*_♂_=–0.33; mean *s*_♀_=0.24; Fig. 3b; Table S10).

**Fig. 3.**
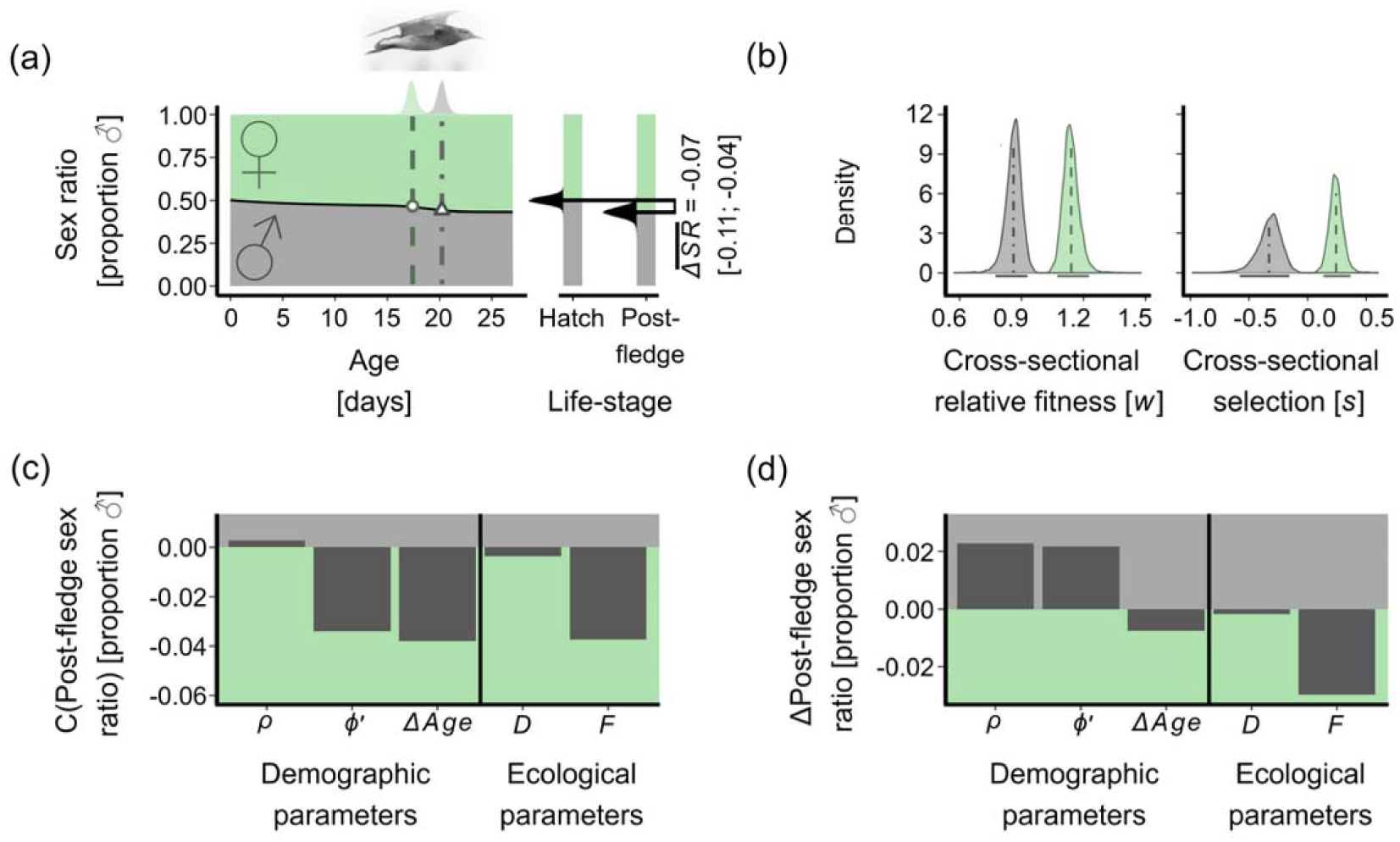
Results of matrix models projecting a female biased post-fledge sex ratio. (a) A single-cohort matrix model predicts a sex ratio (proportion male) decline from hatch to post-fledge. Left, sex ratio declines with cohort age (in days). Shaded areas show population proportions of males (gray) and females (green), black solid line is the mean sex ratio of 10,000 simulations. The mean sex-specific fledging ages are shown as broken lines and shapes (triangles = males; circles = females); the mean sex difference in fledging age across simulations was 2.8 days. This difference is conservative because it is estimated using the posterior probability of fledging at a given age for the model rather than setting a specific fledging age difference for each simulation. Sex-specific fledging age density curves of the simulations are shown on top. Right, sex ratio change (*ΔSR*) between life-stages with mean (horizontal lines) and density curves of simulated estimates for each stage. (b) Simulated density curves of cross-sectional relative fitness *w* and selection coefficients *s* across simulations for males (gray) and females (green). Broken vertical lines indicate the means and solid horizontal lines indicate the 95% credible intervals. (c,d) Results of the life table response experiment showing the contribution of mean parameter estimates to the projected sex ratio decline when the parameter estimate is at its mean value (c), or increased by one standard deviation (d). Demographic parameters: hatching sex ratio *ρ*, cumulative intrinsic survival, and sex difference in fledging age *ΔAge*. Ecological parameters: additive predation mortality *D*, and fledging advantage *F*.

### Life Table Response Experiments

The mean post-fledge SR predicted by the unisex model was higher than the predicted mean post-fledge SR predicted in the two-sex model (Table S10). The predicted mean *ΔSR* between hatching and post-fledging cohort in the unisex model was 53.8% lower than in the two-sex model (Table S10). This indicates that not accounting for intrinsic mortality differences between the sexes has a clear effect on the simulated post-fledge population structure.

The fledging advantage *F* had a strong effect on the SR change meaning that *ΔAge*, which reflected earlier fledging of females than males, had the strongest effect on the SR decline and the negative selection coefficients of males, closely followed by *ϕ* (Fig. 3c; Fig. S9). In contrast, *ρ* and *D* contributed relatively little to the decline in post-fledge SR and to the negative selection coefficients of males (Fig. 3c, Fig. S9a). The sensitivity analysis revealed that the post-fledge SR was most affected by changes in *ρ*, *ϕ′* and *F* whereas the male selection coefficients were only highly affected by changes in *ϕ′* and *F* (Fig. 3d; Fig. S9b).

## Discussion

SR biases have important consequences for sexual selection, sex roles and reproductive rates (Székely *et al*. 2014; Kappeler *et al*. 2023). Yet, when and how SR biases arise remains often unclear. Here we show that in sexually dimorphic Ruffs, sex differences in development and survival during the pre-fledging period result in a female biased SR. Our study provides three major insights into the formation of this bias. First, experimental fledging tests revealed that males and females differ in their fledging age with females fledging 3–5 days earlier than males. Second, this fledging age difference is linked to variation in wing development; the wings of females matured faster than those of males despite males having higher wing growth rates than females for much of the pre-fledging period. Third, the female SR bias increased consistently during the pre-fledging period because males exhibit a higher intrinsic mortality than females. With the onset of fledging the bias increases further because earlier fledging resulted in an additional survival boost for females. Overall, the female bias became clear during the first four weeks post-hatch, about ten months before Ruffs are sexually mature, with fledging age and intrinsic survival differences contributing equally to the observed bias in the fledgling cohort.

### Fledging ages vary considerably among birds

Most of the research to-date has attempted to relate differences between species or populations to the risk of predation (Coslovsky & Richner 2011; Martin *et al*. 2018; de Zwaan *et al*. 2019; Jones *et al*. 2020b). High predation risk should select for faster development, especially in locomotive traits, so that young birds can evade predators sooner and hence increase their survival. In songbirds, high nest predation leads to a shorter nestling period and earlier fledging compared to species with low nest predation (Remeš & Matysioková 2016; Martin *et al*. 2018; Jones *et al*. 2020b). However, early fledging can lead to increased mortality directly after leaving the nest when the flight abilities of newly independent chicks are not yet fully developed (Jones *et al*. 2020b). Resource constraints may lead to trade-offs between investment into growth versus maturation. Rapid maturation should be prioritized over growth when it provides substantial survival benefits. However, when size provides an advantage in adult life, for example, through higher competitiveness during sexual selection, such prioritization of quick maturation may reduce future reproductive success. In Ruffs, fast maturation at the expense of size may be problematic for male Independents, as these males face extremely high mating competition where the top males obtain most of the matings and lower ranking males hardly reproduce (Widemo & Owens 1995; Vervoort & Kempenaers 2020; Tolliver *et al*. 2023). The advantages of large body size in aggressive male-male contests to establish dominance hierarchies on leks may hence have favoured a longer growth period and later maturation in male Independents even if it increases pre-fledgling mortality. In contrast, large body size is not a prerequisite for successful female reproduction, as there is no shortage of male suitors and many females are polyandrous (Lank *et al*. 2002). However, size may still provide a fertility advantage, as large females produce larger eggs resulting in larger chicks, which have improved survival prospects (Blomqvist *et al*. 1997; Giraldo-Deck *et al*. 2022; Eberhart-Hertel *et al*. 2023). Taken together, the divergent sex roles in Ruffs may lead to differently balanced trade-offs in both sexes and serve as an ultimate explanation for the sex differences in pre-sexual mortality and fledging.

A mechanistic explanation for variation in fledging age is based on physical laws. Because of their lower body mass, females require smaller wing areas to generate the required lift-off force than heavier males (Alerstam *et al*. 2007). Consistent with this, Ruff females fledged with shorter wings than males. Intuitively, the positive relationship between body size and fledging age when comparing males and females could indicate stabilizing selection acting on males, where larger males may achieve higher reproductive success than small males but take longer to fledge and therefore may have a higher pre-fledgling mortality risk. Interestingly, within each sex the opposite seemed to be true: large individuals fledged earlier than small ones. This suggests that individuals in good condition were better able to navigate the trade-off between growth and maturation and achieve greater mobility and thus more effective predator evasion earlier than individuals in poorer condition. Wing development at fledging, i.e. the length relative to the fully developed wings in adults, showed little difference between males and females. This suggests that on the path to achieve flight ability wing maturation and growth go hand in hand and females fledged earlier than males because their wings reached the necessary level of maturation at an earlier age.

Another trade-off for precocial chicks concerns the development of different modes of locomotion. For precocial chicks, strong running abilities are essential for their survival as soon as they hatch from their eggs. Chicks run to hunt their invertebrate prey and to escape predators (although for predator evasion, wader chicks heavily rely on camouflage provided by their cryptic plumage early in life (Volkmer *et al*. 2024)). Running speed increases with leg length and well developed leg muscles, whereas for flying, juveniles need to strengthen their pectoralis and supracoideus muscles (Videler 2006). Hence, a trade-off exists between investment into wings and into legs. Even worse, the developing wings must be carried as additional “baggage” and may impede running or reduce jumping performance, unless the wings are used to propel the body in these activities (Heers & Dial 2015).

A bias in the fledgling SR is likely a critical first step towards a biased adult sex ratio (ASR). In waders, biased ASRs often develop from being unbiased at hatching to becoming sex biased before reaching sexual maturity (Eberhart-Phillips *et al*. 2017; Eberhart-Phillips *et al*. 2018). As we show here, the SR bias may originate in the pre-fledging stage, long before sexual maturation. Cross-sectional selection and fitness coefficients suggested that in Ruffs, being a female has a selective advantage at least until fledging, when mortality is typically the highest. Unless the mortality bias is reversed during the consecutive months, males will make up the largest share of the ‘invisible fraction’ (Grafen 1988) that does not reach sexual maturity, as gamete production starts only in the second calendar year. Our population matrix model revealed that the sex differences in fledging age and intrinsic survival contributed about equally to the SR decline with the male proportion dropping from 0.5 at hatching to 0.43 in the fledging population. These results broadly match empirical data on SR in juvenile Ruffs migrating south. Consistent with earlier fledging, juvenile females were captured on average two days earlier than males (Jaatinen *et al*. 2010). Yet, the empirically observed average female SR bias in wild Ruffs was about twice as strong than the average estimate predicted by our model (Jaatinen *et al*. 2010). A potential explanation for this discrepancy is that the parameterization of the model with survival and fledging data derived from chicks raised by hand under benign conditions led to an underestimation of the fledging SR bias. The larger sex is typically more vulnerable to harsh environmental conditions (Benito & González-Solís 2007; Kalmbach *et al*. 2009; Loonstra *et al*. 2019), implying that male offspring mortality is likely higher in natural populations than in captivity. In addition, the use of godwits and lapwings, two species with biparental care may have led to an underestimation of the fledging advantage in Ruffs, where only females provide care to the unfledged young.

ASRs of bird populations are typically male biased (Donald 2007), whereas for Ruffs the fledgling SR bias suggests that the ASR will be female biased. A female-biased ASR has major implications for sexual selection. Mortality costs associated with sexual selection result in the more competitive sex becoming rarer and having a higher reproductive value, i.e., possess a higher future reproductive success, than the more common sex that specializes on care (Jennions & Fromhage 2017; Kappeler 2017), which may have promoted the evolution of the polygamous lekking mating system. Physiological differences between adult males and females might alter the ASR further. Males have higher levels of circulating testosterone than females (Goymann & Wingfield 2014; Loveland *et al*. 2025). Testosterone is important for aggressive dominance contests and it also stimulates somatic growth including growth of bones and muscles (Borski *et al*. 1996; Cox *et al*. 2009). However, high testosterone levels may increase risk taking and suppress immune function (Foo *et al*. 2017; Dunn *et al*. 2024), and hence further reduce long-term survival of males. Females, on the other hand, may have higher mortality risks associated with parental care, as suggested by observations of predation of incubating or brood caring females in our study population (unpublished data). Therefore, further studies on sex differences in adult survival are needed to understand the mortality risks associated with mating competition and parental care and to assess effects on ASR.

In conclusion, we found that differences in development and survival between males and females during early ontogeny shape the juvenile SR in a precocial bird with strong sexual selection. We suggest that in sexually dimorphic species, differences in growth, maturation and fledging age may represent a prominent, but so far neglected, developmental mechanism that results in differential pre-sexual selection acting on males and females. Taken together, our study demonstrates how examining sex differences in fledging age and pre-fledging survival sheds light on how selection acts on the ‘invisible fraction’ of a population (Grafen 1988) and, in turn, may shape demography and sex roles.

## Data accessibility statement

The datasets generated and analysed during the current study as well as the code to generate the presented models, statistics and figures are available in Edmond the Open Research Data Repository of the Max Planck Society, https://doi.org/10.17617/3.AS5DDZ.

## Acknowledgements

We thank the field assistants at Liminganlahti from 2018 until 2023, in particular, Lisa Kreye for their invaluable help. Claudia Scheicher and Petra Neubauer provided animal care of the captive Ruff flock. Melanie Schneider did the molecular sexing. Fränzi Korner-Nievergelt and Susanne Schindler provided advice on the statistical analyses and the population matrix model. We are grateful to the current and past students and members of the Research Group for Behavioural Genetics and Evolutionary Ecology and the Department of Ornithology in Seewiesen, Stephanie Rolio, Aaron Walchuk, Réka, Rosa and Mila Küpper for their help with raising Ruff chicks.

## Statement of authorship

Clemens Küpper, Lina M. Giraldo-Deck and James D. M. Tolliver conceived and designed the study and wrote the first draft. All authors contributed to review and editing. Lina M. Giraldo-Deck and James D. M. Tolliver analysed the data with input from Luke J. Eberhart-Hertel and Clemens Küpper. Kari Koivula, Veli-Matti Pakanen, James D. M. Tolliver, Clemens Küpper, Hanna Algora, Jelena Belojević, Veronika Rohr-Bender and Nelli Rönkä collected data in the field with James D. M. Tolliver and Hanna Algora leading the telemetry. Lina M. Giraldo-Deck, Katrin Martin, Veronika Rohr-Bender, Clemens Küpper, Krisztina Kupán and David B. Lank were responsible for chick raising, measurements and fledging tests in the aviary. Kari Koivula, Veli-Matti Pakanen, Clemens Küpper, David B. Lank and Bart Kempenaers obtained permits for field and aviary work. Clemens Küpper, Kari Koivula, Veli-Matti Pakanen, David B. Lank and Bart Kempenaers supervised the work.

## References

1. Alerstam, T., Rosén, M., Bäckman, J., Ericson, P.G.P. & Hellgren, O. (2007). Flight speeds among bird species: allometric and phylogenetic effects. PLoS Biology, 5, e197.

2. Algora, H., Tolliver, J.D.M., Pakanen, V.-M., Kupán, K., Belojeviæ, J., Rönkä, N. et al. (2025). Nests, Threats, and Leks: Nonrandom Distributions of Nests in Ruffs (Calidris pugnax). Ecology and Evolution, 15, e70997.

3. Anderson, D.J., Reeve, J., Martinez Gomez, J.E., Weathers, W., W., Hutson, S., et al. (1993). Sexual size dimorphism and food requirements of nestling birds. Canadian Journal of Zoology, 71, 2541–2545.

4. Badyaev, A.V. (2002). Growing apart: an ontogenetic perspective on the evolution of sexual size dimorphism. Trends in Ecology & Evolution, 17, 369–378.

5. Benito, M.M. & González-Solís, J. (2007). Sex ratio, sex-specific chick mortality and sexual size dimorphism in birds. Journal of Evolutionary Biology, 20, 1522–1530.

6. Blanckenhorn, W.U., Dixon, Anthony F.G., Fairbairn, Daphne J., Foellmer, Matthias W., Gibert, P., Linde, Kim van d., et al. (2007). Proximate causes of Rensch’s rule: Does sexual size dimorphism in arthropods result from sex differences in development time? The American Naturalist, 169, 245–257.

7. Blomqvist, D., Johansson, O.C. & Götmark, F. (1997). Parental quality and egg size affect chick survival in a precocial bird, the lapwing Vanellus vanellus. Oecologia, 110, 18–24.

8. Boomsma, J.J. & Grafen, A. (1990). Intraspecific variation in ant sex ratios and the Trivers-Hare hypothesis. Evolution, 44, 1026–1034.

9. Borski, R.J., Tsai, W., DeMott-Friberg, R. & Barkan, A.L. (1996). Regulation of somatic growth and the somatotropic axis by gonadal steroids: primary effect on insulin-like growth factor I gene expression and secretion. Endocrinology, 137, 3253–3259.

10. Caswell, H. (2001). Matrix Population Models: Construction, Analysis, and Interpretation. Second edn. Oxford University Press, Sinauer, Sunderland, MA.

11. Chiba, S., Iwamoto, A., Shimabukuro, S., Matsumoto, H. & Inoue, K. (2023). Mechanisms that can cause population decline under heavily skewed male-biased adult sex ratios. Journal of Animal Ecology, 92, 1893–1903.

12. Clutton-Brock, T.H., Albon, S.D. & Guinness, F.E. (1985). Parental investment and sex differences in juvenile mortality in birds and mammals. Nature, 313, 131–133.

13. Coslovsky, M. & Richner, H. (2011). Predation risk affects offspring growth via maternal effects. Functional Ecology, 25, 878–888.

14. Cox, R.M., Stenquist, D.S. & Calsbeek, R. (2009). Testosterone, growth and the evolution of sexual size dimorphism. Journal of Evolutionary Biology, 22, 1586–1598.

15. Darwin, C. (1871). Descent of man, and selection in relation to sex. J. Murray, London.

16. de Zwaan, D.R., Camfield, A.F., MacDonald, E.C. & Martin, K. (2019). Variation in offspring development is driven more by weather and maternal condition than predation risk. Functional Ecology, 33, 447–456.

17. Dunn, S.E., Perry, W.A. & Klein, S.L. (2024). Mechanisms and consequences of sex differences in immune responses. Nature Reviews Nephrology, 20, 37–55.

18. Düsing, C. (1884). Die Regulierung des Geschlechtsverhätnisses bei der Vermehrung der Menschen, Tiere und Pflanzen. Jenaische Zeitschrift für Naturwissenschaft, 17, 593–940.

19. Eberhart-Hertel, L., Safari, I., Makomba, P., Hertel, A. & Goymann, W. (2024). Early-life demographic processes do not drive adult sex ratio biases and mating systems in sympatric coucal species. Functional Ecology, 38, 1779 1795.

20. Eberhart-Hertel, L.J., Rodrigues, L.F., Krietsch, J., Hertel, A.G., Cruz-López, M., Vázquez-Rojas, K.A. et al. (2023). Egg size variation in the context of polyandry: a case study using long-term field data from snowy plovers. Evolution, 77, 2590–2605.

21. Eberhart-Phillips, L.J., Küpper, C., Carmona-Isunza, M.C., Vincze, O., Zefania, S., Cruz-López, M. et al. (2018). Demographic causes of adult sex ratio variation and their consequences for parental cooperation. Nature Communications, 9, 1651.

22. Eberhart-Phillips, L.J., Küpper, C., Miller, T.E.X., Cruz-López, M., Maher, K.H., dos Remedios, N., et al. (2017). Sex-specific early survival drives adult sex ratio bias in snowy plovers and impacts mating system and population growth. Proceedings of the National Academy of Sciences, 114, E5474

23. .Fisher, R.A. (1930). The genetical theory of natural selection. Clarendon Press, Oxford, England.

24. Foo, Y.Z., Nakagawa, S., Rhodes, G. & Simmons, L.W. (2017). The effects of sex hormones on immune function: a meta-analysis. Biological Reviews, 92, 551–571.

25. Gelman, A. & Su, Y. (2015). arm: Data analysis using regression and multilevel/hierarchical models [Online]. R package version 1.8–6.

26. Giraldo-Deck, L.M., Goymann, W., Safari, I., Dawson, D.A., Stocks, M., Burke, T. et al. (2020). Development of intraspecific size variation in black coucals, white-browed coucals and ruffs from hatching to fledging. Journal of Avian Biology, 51.

27. Giraldo-Deck, L.M., Loveland, J., Goymann, W., Tschirren, B., Burke, T., Kempenaers, B. et al. (2022). Intralocus conflicts associated with a supergene. Nature Communications, 13, 1384.

28. Goymann, W. & Wingfield, J.C. (2014). Male-to-female testosterone ratios, dimorphism, and life history—what does it really tell us? Behavioral Ecology, 25, 685–699.

29. Grafen, A. (1988). On the uses of data on lifetime reproductive success. In: Reproductive success. Studies of individual variation in contrasting breeding systems (ed. Clutton-Brock, T). University of Chicago Press Chicago, IL, pp. 454–471.

30. Heers, A.M. & Dial, K.P. (2015). Wings versus legs in the avian bauplan: development and evolution of alternative locomotor strategies. Evolution, 69, 305–320.

31. Heinsohn, R., Olah, G., Webb, M., Peakall, R. & Stojanovic, D. (2019). Sex ratio bias and shared paternity reduce individual fitness and population viability in a critically endangered parrot. Journal of Animal Ecology, 88, 502–510.

32. Hirst, A.G., Bonnet, D., Conway, D.V.P. & Kiørboe, T. (2010). Does predation controls adult sex ratios and longevities in marine pelagic copepods? Limnology and Oceanography, 55, 2193–2206.

33. Hogan-Warburg, A.J. (1966). Social behavior of the ruff Philomachus pugnax L. Ardea, 54, 109–229.

34. Hugie, D.M. & Lank, D.B. (1997). The resident’s dilemma: a female choice model for the evolution of alternative mating strategies in lekking male ruffs (Philomachus pugnax). Behavioral Ecology, 8, 218–225.

35. Jaatinen, K., Lehikoinen, A. & Lank, D.B. (2010). Female-biased sex ratios and the proportion of cryptic male morphs of migrant juvenile ruffs (Philomachus pugnax) in Finland. Ornis Fennica, 87, 125–134.

36. Jennions, M.D. & Fromhage, L. (2017). Not all sex ratios are equal: the Fisher condition, parental care and sexual selection. Philosophical Transactions of the Royal Society B: Biological Sciences, 372, 20160312.

37. Jones, T.M., Benson, T.J. & Ward, M.P. (2020a). Does the size and developmental stage of traits at fledging reflect juvenile flight ability among songbirds? Functional Ecology, 34, 799–810.

38. Jones, T.M., Brawn, J.D., Ausprey, I.J., Vitz, A.C., Rodewald, A.D., Raybuck, D.W. et al. (2020b). Parental benefits and offspring costs reflect parent–offspring conflict over the age of fledging among songbirds. Proceedings of the National Academy of Sciences, 117, 30539–30546.

39. Jukema, J. & Piersma, T. (2006). Permanent female mimics in a lekking shorebird. Biology Letters, 2, 161–164.

40. Kalmbach, E., Furness, R.W. & Griffiths, R. (2005). Sex-biased environmental sensitivity: natural and experimental evidence from a bird species with larger females. Behavioral Ecology, 16, 442–449.

41. Kalmbach, E., Griffiths, R. & Furness, R.W. (2009). Sex-specific growth patterns and effects of hatching condition on growth in the reversed sexually size-dimorphic great skua Stercorarius skua. Journal of Avian Biology, 40, 358–368.

42. Kappeler, P.M. (2017). Sex roles and adult sex ratios: insights from mammalian biology and consequences for primate behaviour. Philosophical Transactions of the Royal Society B: Biological Sciences, 372, 20160321.

43. Kappeler, P.M., Benhaiem, S., Fichtel, C., Fromhage, L., Höner, O.P., Jennions, M.D. et al. (2023). Sex roles and sex ratios in animals. Biological Reviews, 98, 462–480.

44. Koivula, K., Algora, H., Airaksinen, E., Belojeviæ, J., Küpper, C., Oranen, M. et al. (2025). Increased wind flood frequency leads to decreased nest success of endangered waders in managed shore meadows. Biological Conservation, 302, 110970.

45. Kokko, H. & Jennions, M.D. (2008). Parental investment, sexual selection and sex ratios. Journal of Evolutionary Biology, 21, 919–948.

46. Krackow, S. (2002). Why Parental Sex Ratio Manipulation is Rare in Higher Vertebrates (Invited Article). Ethology, 108, 1041–1056.

47. Küpper, C., Stocks, M., Risse, J.E., dos Remedios, N., Farrell, L.L., McRae, S.B., et al. (2016). A supergene determines highly divergent male reproductive morphs in the ruff. Nature Genetics, 48, 79–83.

48. Lamichhaney, S., Fan, G., Widemo, F., Gunnarsson, U., Thalmann, D.S., Hoeppner, M.P. et al. (2016). Structural genomic changes underlie alternative reproductive strategies in the ruff (Philomachus pugnax). Nature Genetics, 48, 84–88.

49. Lank, D.B., Farrell, L.L., Burke, T., Piersma, T. & B., M.S. (2013). A dominant allele controls development into female mimic male and diminutive female ruffs. Biology Letters, 9, 20130653.

50. Lank, D.B., Smith, C.M., Hanotte, O., Ohtonen, A., Bailey, S. & Burke, T. (2002). High frequency of polyandry in a lek mating system. Behavioral Ecology, 13, 209–215.

51. Le Galliard, J.-F., Fitze, P.S., Ferrière, R. & Clobert, J. (2005). Sex ratio bias, male aggression, and population collapse in lizards. Proceedings of the National Academy of Sciences, 102, 18231–18236.

52. Liker, A., Freckleton, R.P. & Székely, T. (2013). The evolution of sex roles in birds is related to adult sex ratio. Nature Communications, 4, 1587–1587.

53. Linnen, C. & Hoekstra, H.E. (2009). Measuring natural selection on genotypes and phenotypes in the wild. In: Cold Spring Harbor Sumposia on Quantitative Biology. Cold Spring Harbor Labratory Press Cold Spring Harbor, pp. 155–168.

54. Loonstra, A.H.J., Verhoeven, M.A., Senner, N.R., Hooijmeijer, J.C.E.W., Piersma, T. & Kentie, R. (2019). Natal habitat and sex-specific survival rates result in a male-biased adult sex ratio. Behavioral Ecology, 30, 843–851.

55. Loveland, J.L., Zemella, A., Jovanoviæ, V.M., Möller, G., Sager, C.P., Bastos, B. et al. (2025). A single gene orchestrates androgen variation underlying male mating morphs in ruffs. Science, 387, 406–412.

56. Lovich, J.E. & Gibbons, J.W. (1990). Age at maturity influences adult sex ratio in the turtle Malaclemys terrapin. Oikos, 59, 126–134.

57. Martin, T.E., Tobalske, B., Riordan, M.M., Case, S.B. & Dial, K.P. (2018). Age and performance at fledging are a cause and consequence of juvenile mortality between life stages. Science Advances, 4, eaar1988.

58. Mason, L.R., Smart, J. & Drewitt, A.L. (2018). Tracking day and night provides insights into the relative importance of different wader chick predators. Ibis, 160, 71–88.

59. Nijhout, H.F., Roff, D.A. & Davidowitz, G. (2010). Conflicting processes in the evolution of body size and development time. Philosophical Transactions of the Royal Society B: Biological Sciences, 365, 567–575.

60. Oakes, E.J. (1992). Lekking and the evolution of sexual dimorphism in birds: comparative approaches. The American Naturalist, 140, 665–684.

61. Parker, G.A. (1992). The evolution of sexual size dimorphism in fish. Journal of Fish Biology, 41, 1–20.

62. Promislow, D.E.L. (1992). Costs of sexual selection in natural populations of mammals. Proc. R. Soc. Lond. B. Biol. Sci., 247, 203–210.

63. R Development Core Team (2024). R: A language and environment for statistical computing. R Foundation for Statistical Computing Vienna, Austria.

64. Regan, C.E., Medill, S.A., Poissant, J. & McLoughlin, P.D. (2020). Causes and consequences of an unusually male-biased adult sex ratio in an unmanaged feral horse population. Journal of Animal Ecology, 89, 2909–2921.

65. Remeš, V. & Matysioková, B. (2016). Survival to independence in relation to pre-fledging development and latitude in songbirds across the globe. Journal of Avian Biology, 47, 610–618.

66. Richner, H. (1991). The Growth Dynamics of Sexually Dimorphic Birds and Fisher’s Sex Ratio Theory: Does Sex-Specific Growth Contribute to Balanced Sex Ratios? Functional Ecology, 5, 19–28.

67. Ricklefs, R.E. (1979). Adaptation, constraint, and compromise in avian postnatal development. Biological Reviews, 54, 269–290.

68. Schekkerman, H., Teunissen, W. & Oosterveld, E. (2009). Mortality of black-tailed godwit Limosa limosa and Northern lapwing Vanellus vanellus chicks in wet grasslands: influence of predation and agriculture. Journal of Ornithology, 150, 133–145.

69. Schindler, S., Gaillard, J.M., Grüning, A., Neuhaus, P., Traill, L.W., Tuljapurkar, S. et al. (2015). Sex[specific demography and generalization of the Trivers–Willard theory. Nature, 526, 249–252.

70. Staufer, M., Burgstaller, S., Horvath, A. & Landler, L. (2023). Temporal and spatial variations in local sex ratios in a suburban population of the European green toad Bufotes viridis. BMC Ecology and Evolution, 23, 6.

71. Svensson, E.I., Willink, B., Duryea, M.C. & Lancaster, L. (2019). Temperature drives pre-reproductive selection and shapes the biogeography of a female polymorphism. Ecology Letters.

72. Székely, T., Liker, A., Freckleton, R.P., Fichtel, C. & Kappeler, P.M. (2014). Sex-biased survival predicts adult sex ratio variation in wild birds. Proceedings of the Royal Society B: Biological Sciences, 281, 20140342.

73. Tolliver, J.D.M., Kupán, K., Lank, D.B., Schindler, S. & Küpper, C. (2023). Fitness benefits from co-display favour subdominant male–male partnerships between phenotypes. Animal Behaviour, 197, 131–154.

74. Torres, R. & Drummond, H. (1997). Female-biased mortality in nestlings of a bird with size dimorphism. Journal of Animal Ecology, 66, 859–865.

75. Trivers, R.L. & Willard, D.E. (1973). Natural Selection of parental ability to vary the sex ratio of offspring. Science, 179, 90–92.

76. Umlauf, N., Klein, N. & Zeileis, A. (2018). BAMLSS: Bayesian additive models for location, scale, and shape (and beyond). Journal of Computational and Graphical Statistics, 27, 612–627.

77. van Rhijn, J.G. (1991). The ruff. Poyser, London.

78. Veran, S. & Beissinger, S.R. (2009). Demographic origins of skewed operational and adult sex ratios: perturbation analyses of two[sex models. Ecology Letters, 12, 129–143.

79. Vervoort, R. & Kempenaers, B. (2020). Variation in lek attendance and copulation success of Independent and Satellite male ruffs Calidris pugnax. Ardea, 107, 303–320, 318.

80. Videler, J.J. (2006). Avian flight. Oxford University Press, Oxford.

81. Volkmer, T., Kupán, K., Rohr-Bender, V.A., Guirao-Ortiz, M., Cruz-López, M., del Angel, S.G. et al. (2024). Hidden in plain sight: camouflage and hiding behaviour of wild precocial chicks in an open landscape. Behavioral Ecology and Sociobiology, 78, 73.

82. West, S.A. & Sheldon, B.C. (2002). Constraints in the Evolution of Sex Ratio Adjustment. Science, 295, 1685–1688.

83. Widemo, F. (1998). Alternative reproductive strategies in the ruff Philomachus pugnax: a mixed ESS? Animal Behaviour, 56, 329–336.

84. Widemo, F. & Owens, I.P.F. (1995). Lek size, male mating skew and the evolution of lekking. Nature, 373, 148–151.

