## Supplement information for "Sex differences in development alter the fledgling sex ratio in a lekking bird with strong sexual size dimorphism"

### Supplementary Methods

*Rearing and measurements of captive chicks.* In both locations, Burnaby (Canada) and Seewiesen (Germany) we collected eggs several times daily during daylight hours. We incubated the eggs at 37.5°C and 55% humidity until hatching or death of the embryos. After hatching, we collected the remaining blood vessels from the inside of the eggshell, or took a small blood sample from the chick for genetic sex and morph identification using single nucleotide markers (Giraldo-Deck *et al.* 2020). To control for potential maternal effects, we assigned mothers to each chick through genetic parentage analyses using polymorphic microsatellite markers (Giraldo-Deck *et al.* 2022). Hatched chicks were individually colour-ringed and hand-raised with other chicks of similar age in heterosexual groups under *ad libitum* food until fledging. Shortly after fledging, they were released to the adult flock. We measured body mass and tarsus length during their first or second winter as adults as indicators of their adult body size. To relate the development of flight ability, wing growth, and feather maturation to individual adult body size, we weighed the birds and measured tarsus length in November or December of their first or second year. This body mass represents the asymptotic lean adult body mass in Ruffs.

*Wing, feather and body size measurements.* To compare wing growth between sexes we measured wing length (to the nearest mm) as ‘unflattened wing’ (Evans 1986) in chicks starting on day 5 post-hatch, two to three times per week. For logistical reasons, measurement schedules slightly varied between breeding seasons. In 2019, we took daily measurements from day 16 post-hatch until the bird had fledged. From 2020 until 2023, we measured wing length only every third day and on the day of fledging. To evaluate variation in feather maturation, we determined feather emergence of the second outmost primary, which is the longest wing feather, whenever we measured the wing length. To obtain the proportion of the mature, unfurled region of the feather, we measured its total length from the papilla to the tip and its emerged length from the most distal part of the sheath to the tip.

To compare wing length and wing growth between sexes we first modelled posterior means of wing length and their 95% credible intervals (CrIs) in relation to age for each sex. For this, we used the R package BAMLSS, a flexible Bayesian generalized additive model framework (GAMM) based on Markov chain Monte Carlo simulations that can include random factors (Umlauf *et al.* 2018). We modelled each sex separately because fixed effects cannot be specified in BAMLSS and our main interest was to examine differences in the shape of growth curves between males and females. For each model, we z-transformed the variables wing length and age by setting their means to zero and their standard deviation to one. We included ‘mother ID’, ‘Individual ID’ and an interaction between ‘Individual ID’ and ‘age’ as a random factor in all models to allow for individual specific growth curves. In separate models, we estimated posterior means and 95% CrI of wing growth rates and of the proportion of adult wing length in relation to age. We calculated growth rate for each individual as:

growth rate = (*Wt* - *Ws*) / (*t* - *s*)

where *Wt* is the wing length at age *t* and *Ws* is the preceding wing length to *Wt* at age *s*. We then calculated the proportion of adult wing length by dividing *Wt* with the adult wing length *WAd*.

To examine feather maturation we compared feather emergence, i.e. the length of the emerged unfurled feather section of the second outmost primary between sexes. We used the same GAMM structure as for wing growth described above and z-transformed values for variables feather emergence and age. In separate models, we evaluated both the rate and proportion of feather emergence in relation to age. We calculated rate and proportion of feather emergence as:

rate of feather emergence = (*Ft* - *Fs*) / (*t* - *s*)

where *Ft* is the length of the emerged part of the second outermost primary at age *t* and *Fs* is the corresponding length at age *s*. We calculated the proportion of feather emergence by dividing *Ft* with the total length (papilla to tip) of the second outmost primary.

*Chick survival.* We collected survival data of wild chicks by opportunistic re-captures of ringed chicks and radio tracking of tagged chicks from 2019 until 2022. We used two types of radio transmitters, VHF and UHF tags, that we glued to kinesiology tape, which we then attached to the pelvic girdle of chicks with water-based latex adhesive (Copydex) (Mason *et al.* 2017). In line with the 5% of body mass rule (Grant & Oring 1997; Lees *et al.* 2019), we used 0.5 g VHF tags (manufacturers: Telemetrie-Service Dessau, Germany and Holohil, Canada) on small chicks (body mass: 10–13 g), and 0.6–0.7 g UHF tags (manufacturer: Cellular Tracking Technologies (CTT), USA) on large chicks (body mass: >14 g). We then returned chicks to their nest and checked the nest once more within 24 h to establish that the chicks had left the nest with the mother. We typically marked only two chicks to minimize brood effects on survival estimates, except for seven broods where we marked three to four chicks. We then monitored radio tagged chicks by either tracking them on foot approximately every second day with handheld radio receivers attached to Yagi antenna, and/or using a grid of automated radio receivers which transmitted data to a centralized receiving station. We followed chicks until we could not locate them after several consecutive tracking attempts, found them dead or we confirmed predation by abrupt changes in the tracking data (Fig. S3.

To quantify sex- and age-specific intrinsic survival and fledging probabilities, we collected data from the captive Ruff population, described above, from 2018 to 2023 (Table S2). We monitored chick survival from hatching until 27 days of age. By this age, all juveniles of both sexes had either fledged or died. For the survival data we used only Independent chicks that had either Independent or Satellite mothers because chicks of Faeder mothers have higher mortality (Giraldo-Deck *et al.* 2022). We used mixed-effects known fate survival models (Korner-Nievergelt *et al.* 2024) to estimate posterior chick intrinsic survival and fledging probabilities (Figs. S4,S5). The intrinsic survival and fledging models had the same fixed effects structure with ‘Age’ (in days and z-transformed), ‘Sex’, and the interaction of Age and Sex as predictors. However, the interaction between Age and Sex only had a clear relationship in the fledging model and was therefore not included in the final survival model (Tables S4, S5). In terms of random effects, the fledging model included both Mother ID (‘Mother’) and Cohort year (‘Year’) whereas the survival model only had Year (Tables S4, S5). We coded for chick survival as ‘1’ (survived) and ‘0’ (dead), within the survival model. For the fledging model, the probability of not flying was coded ‘1’ and flying coded as ‘0’ for the response variable. We bounded ‘Age’ in the fledging model by the empirical observations in our fledging tests (min age=16; max age=27).

To quantify additive predation mortality *D*, we prepared a second known fate survival model to estimate wild survival. As the sample size of tracked chicks was limited (82 chicks from 38 clutches), we did not fit sex as predictor. We used ‘Cohort ID’ and ‘Brood ID’ as random effects. Because our tracking was limited to chicks up to an age of 11 days, our model had a lower age range (0 -11) than our intrinsic survival model (0–27 days). The model showed no clear effect of Age on survival and hence we used the mean posterior distribution as wild survival (Table S6). We derived *D* from parameters we were able to estimate from empirical data: the survival of chicks in the wild $\phi_{w}$, and the survival of chicks in the aviary (intrinsic survival) $\phi_{I}$. Because we censored out chicks from the wild that died from causes other than predation (e.g., cattle trampling, drowning, found dead with no clear cause etc.), we assumed that $\phi_{w}$ was the product of chicks that survived from intrinsic sources of mortality and predation sources of mortality, summarized in the following equation:

$\phi_{w}$= $\phi_{I}\phi_{P}$

where $\phi_{I}$ and $\phi_{P}$ are the survival of chicks from intrinsic mortality sources and predation mortality sources, respectively. From this equation, we substituted 1 – *D* for $\phi_{P}$ where *D* is the proportion of chicks that die from additive predation.

$\phi_{w}$= $\phi_{I} (1-D)$

We then solved for *D*,

$\frac{\phi_{w}}{\phi_{I}}$ = $1-D$

$\frac{\phi_{w}}{\phi_{I}}$ $-$1 = $- D$

$D$ = 1 $-$ $\frac{\phi_{w}}{\phi_{I}}$

and obtained the posterior of *D* from posteriors of $\phi_{w}$and $\phi_{I}$. We removed negative values from the posterior distribution because they are not biologically plausible.

*Fledging advantage.* Fledging enables chicks to evade ground predators and, hence, boosts their survival. Comparing the survival of radio-tracked wild and captive chicks we quantified the additive predation mortality that Ruff chicks experience in the wild. Because the radio tracking ended before the fledgling period, we estimated the fledging survival boost in our model using supplementary data from two wader species that use the same breeding habitat as Ruffs (Schekkerman *et al.* 2009). We derived the fledging advantage by taking the product of the daily survival estimates before and after the age at first fledging for Black-tailed Godwits (*Limosa limosa*; fledge at day 25) and Northern Lapwings (*Vanellus vanellus*; fledge at day 33) (Fig. S6). We determined the period-specific daily survival rates for each species by taking the nth root of the cumulative survival for pre-fledging and fledging periods, where n was the number of days in each period. Taking the nth root allowed us to account for the difference in the span of ages that godwits and lapwings spent in the pre- and post-fledge stages. For Ruffs, we used the mean difference between pre-fledge daily survival and post-fledge daily survival from godwits and lapwings as fledging advantage in our model. We consider our derived fledging advantage to be a conservative estimate, because both godwits and lapwings exhibit biparental care until fledging, whereas in Ruffs only the female cares and is able to defend the unfledged chicks.

*Matrix model.* For HSR, survival, and fledging probability, we used four chains of 3,500 simulations, with a 1,000-sample warm-up, to obtain a total of 10,000 samples from the posterior distributions. Model selection involved two steps. First, we estimated a global model with predictive parameters of biological relevance. Second, we reduced the complexity of the model by only including parameters that clearly differed from zero (Dushoff *et al.* 2019). We projected the population through an array of two-sex, age-specific matrices, whereby each matrix represented a chick’s transition from a given day of age to the next.

The general projection for each matrix per time step *t* is given by:

$$m_{t+1}= M_{t}m_{t}$$

where ***m*** is a 4 $\times$ 1 vector of the age- and sex-specific population distributed across two life-stages (chick = *C* and juvenile = *J*):

$\boldsymbol{m}_{\boldsymbol{t}}$= $\left[ \begin{matrix} n_{t♀}^{C} \\ n_{t♂}^{C} \\ n_{t♀}^{J} \\ n_{t♂}^{J} \end{matrix} \right]$

and $\boldsymbol{M}$(*t*) is a 4 $\times$ 4 matrix of age- and sex-specific transition probabilities:

$$\left[ \begin{matrix} \phi_{t♀}(1- D) (1-\psi_{t♀})\text{ } & 0 & 0 & 0 \\ 0 & \phi_{t♂} (1- D)(1-\psi_{t♂})\text{ } & 0 & 0 \\ \phi_{t♀} (1- D)\text{ }\psi_{t♀} & 0 & \phi_{t♀} (1-\left( D - F \right)) & 0 \\ 0 & \phi_{t♂} (1-\left( D - F \right))\text{ }\psi_{t♂} & 0 & \phi_{t♂} (1-(D - F)) \end{matrix} \right]$$

Here age-specific *t* transition probabilities are intrinsic survival *ϕ* and the probability of a chick fledging $\psi$. Chicks remained in the chick stage at a rate equal to $1-\psi_{tS}$ and survived at a rate equal to $\phi_{tS} (1-D)$, where *D* is the additive predation mortality. Chicks fledged and became juveniles at a rate equal to $\psi_{tS}$ and juveniles survived at a rate equal to $\phi_{tS}(1-(D - F)$), where *F* is the fledging advantage.

*Estimating sensitivities in life table response experiment.* We estimated sensitivities using numerical methods that independently perturbed our input parameters within their posterior distributions. Each parameter was perturbed 1,000 times with one cohort projected from each perturbation. From these perturbations, we estimated parameter-specific splines and derived $\frac{\partial y}{\partial\theta}$ from each spline (Eberhart-Phillips *et al.* 2017), where $\frac{\partial y}{\partial\theta}$ is the sensitivity of the response variable *y* (i.e., in our case, the post-fledging sex ratio) to the observed mean vital rate $\theta$ in the two-sex model. Since *F* had no posterior distribution, we sampled from the difference in survival from the pre-fledge and post-fledge survival estimates derived from (Schekkerman *et al.* 2009). The sensitivity analysis required the same number of predictor values as response values to estimate the parameter specific-spline. To estimate sensitivities of post-fledge SR and the male selection coefficient, *s_♂_*, we collapsed daily estimates of *ϕ* and *ψ* for each simulation into a single estimate. For *ϕ,* we took the cumulative intrinsic survival, $\phi^{'}$, whereas for *ψ,* we used the mean difference in fledging age, Δ*Age*.

The formula to calculate the theoretical unit change in the post-fledge SR (y) and *s_♂_* that we used was:

$\Delta y\left( {SD}_{\theta} \right)= \frac{\partial y}{\partial\theta}{SD}_{\theta}$

where $\theta$ is the parameter of interest, $y$ is the response variable of interest, and $\frac{\partial y}{\partial\theta}$ is the sensitivity of $y$ to $\theta$ in the two-sex model, and *SD* is the standard deviation of the distribution of $\theta$.

To calculate the individual contributions of demographic and ecological parameters to the fledging sex ratio, we decomposed the demographic parameters (ρ,$\phi^{'}$, and 𝛥𝐴𝑔𝑒) and ecological parameters (*F* and *D*) into their lower-level sex-biases and, for ecological parameters, their differences from zero to calculate their relative contributions to the post-fledge SR and male selection coefficients in our two-sex model. These demographic parameter contributions *C*, were obtained by the formula:

$C_{y}$(θ) = ($\theta_{r}$ –$\theta_{t}$)$\frac{\partial y^{'}}{\partial\theta^{'}}$

Here $\frac{\partial y^{'}}{\partial\theta^{'}}$ is the sensitivity of the response variable *y* to perturbations in the parameter 𝜃 in the array of matrices 𝐌′ (𝑡 = 0 … 27). For calculating *C* for the demographic parameters, $\theta_{r}$ was the male mean parameter estimate across the two-sex model simulations (i.e. male survival or male fledging age) and $\theta_{t}$ was the mean female parameter estimate across those same simulations. For the ecological parameters, $\theta_{r}$ was the mean parameter estimate across simulations and $\theta_{t}$ was set at zero. Finally, ρ, the hatching sex ratio, was set at 0.5.

### Supplementary Tables and Figures

**Table S1. Sample sizes and sex information of wild Ruff chicks sampled at hatching at Pitkänokka meadows, Liminka Bay, Finland from 2016 until 2023. The data were used to inform the Population Matrix Model.** Shown in the table are the number of chicks (♀: female chicks; ♂: male chicks) and the number of clutches sampled per year.

| **Years** | **Chicks** | **Clutches** | **Sex** | |
| --- | --- | --- | --- | --- |
|  |  |  | **♀** | **♂** |
| 2016 | 55 | 18 | 27 | 28 |
| 2017 | 87 | 37 | 41 | 46 |
| 2018 | 150 | 52 | 73 | 77 |
| 2019 | 76 | 31 | 43 | 33 |
| 2020 | 51 | 19 | 25 | 26 |
| 2021 | 59 | 19 | 29 | 30 |
| 2022 | 70 | 24 | 33 | 37 |
| 2023 | 108 | 42 | 56 | 52 |
| All years | 656 | 242 | 327 | 329 |

**Table S2. Sample sizes, sex and fates of captive Ruff chicks used to model intrinsic survival and fledging age.** Alive chicks survived at least 27 days after hatching, Dead chicks survived less than 27 days after hatching and Censored chicks were used for other experiments that involved tissue sampling before 27 days after hatching and were hence censored in the survival models. Note that all chicks that were tested for fledging survived until they fledged. Hence, alive, died, and censor are filled with ‘—’.

| **Analysis** |  | **Total chicks** | **Fate** | | | **Sex** | |
| --- | --- | --- | --- | --- | --- | --- | --- |
|  |  |  | **Alive** | **Dead** | **Censored** | **♀** | **♂** |
| Survival |  | 377 | 271 | 74 | 32 | 170 | 207 |
| Fledging |  | 196 | — | — | — | 108 | 88 |

**Table S3.** **Posterior distribution of parameter estimates of potential predictors on hatching sex ratio (i.e., probability of a chick being male)**. The *Initial model* included a subset of our sampled chicks, *n* = 486 from 173 clutches collected over 8 years (2016 – 2023), because we did not have nest initiation data for all clutches. The model examined whether the relative nest initiation date (Season) influenced the hatching sex ratio (HSR) in a given clutch per a given year. The model included Season (scaled to a mean of zero and standard deviation of 1) as linear and quadratic fixed effects and Clutch ID and Cohort ID as random effects. Since there was no clear effect of Season or Season^2^, we used the *Selected Model* that did not include seasonal effects to inform the matrix model. This *Selected model* was estimated from our whole data set, 656 chicks from 242 clutches, because seasonal data did not limit this model’s sample size. The table provides the parameter estimates that are not back-transformed from the logit-link function. β_Intercept_ provides the estimated HSR for the mean nest initiation date (Julian date 146/Gregorian date 26 May). β_Season_ and β_Season_^2^ provide the estimated changes in HSR per standard deviation of age and squared standard deviation of age, respectively. σ_Clutch ID_ and σ_Cohort ID_ indicate the estimates for the between-clutch standard deviation and the between-cohort standard deviation, respectively. Residual standard deviations were 0.32 (0.31 – 0.32) and 0.33 (0.33 – 0.33) for the *Initial Model* and the *Selected Model*, respectively. The table contains the mean, the 95% CrI (2.5% and the 97.5% quantiles) and the posterior probability of the hypothesis that the parameter, β, is smaller than zero. Probabilities in bold are comparable to a significant effect (P<0.05) according to frequentist statistics.

| **Hatching sex ratio models** | | | | |
| --- | --- | --- | --- | --- |
| **Parameter** | mean | 2.5% | 97.5% | P (β < 0) |
| ***Initial model*** | | | | |
| **β_Intercept_** | -0.125 | -0.39 | 0.134 | 0.838 |
| **β_Season_** | -0.052 | -0.29 | 0.188 | 0.664 |
| **β_Season_^2^** | 0.031 | -0.09 | 0.153 | 0.306 |
| $\boldsymbol{\sigma}$**_Clutch ID_** | 0.031 | < 0.001 | 0.163 | — |
| $\boldsymbol{\sigma}$**_Cohort ID_** | 0.04 | < 0.001 | 0.217 | — |
| ***Selected model*** | | | | |
| **β_Intercept_** | 0.006 | -0.171 | 0.188 | 0.473 |
| $\boldsymbol{\sigma}$**_Clutch ID_** | 0.027 | < 0.001 | 0.138 | — |
| $\boldsymbol{\sigma}$**_Cohort ID_** | 0.016 | < 0.001 | 0.096 | — |

**Table S4.** **Posterior distribution of parameter estimates for potential predictors on intrinsic survival (daily chick survival probability from hatching until the age of 27 days from our captive population)**. The *Initial model* included chick age (scaled to a mean of zero and standard deviation of 1), sex and their interaction as fixed effects and Cohort ID as random factor. Because there was no clear interaction effect, our *Selected model* for informing the matrix model included only the fixed effects of chick age and sex, and the random effect of cohort. The table provides the parameter estimates that are not back-transformed from the logit-link function. β_Intercept_ provides the estimated intrinsic survival for a male chick at an age of 13.5 days (i.e., the mean of the set 0 to 27 days). β_Age_ provides the estimated change in intrinsic survival per standard deviation of age, β_Female_ provides the estimated difference in intrinsic survival between female and male individuals at an age of 13.5 days, and β_Age:Female_ provides the estimated interaction between chick age and sex. σ_Cohort ID_ indicates the estimate for the between-cohort standard deviation. In both models the residual standard deviations were similar 0.01 (0.01 – 0.02). The table contains the mean, the 95% CrI (2.5% and 97.5% quantiles) and the posterior probability of the hypothesis that the parameter, β, is smaller than zero. Probabilities in bold are comparable to a significant effect (P<0.05) according to frequentist statistics.

| **Intrinsic survival models** | | | | |
| --- | --- | --- | --- | --- |
| **Parameter** | mean | 2.5% | 97.5% | P (β < 0) |
| ***Initial model*** | | | | |
| **β_Intercept_** | 5.159 | 4.367 | 5.975 | **< 0.001** |
| **β_Age_** | 1.235 | 0.835 | 1.663 | **< 0.001** |
| **β_Female_** | 0.471 | -0.238 | 1.189 | 0.093 |
| **β_Age:Female_** | -0.294 | -0.907 | 0.312 | 0.831 |
| $\boldsymbol{\sigma}$**_Cohort ID_** | 0.165 | 0.07 | 0.346 | — |
| ***Selected model*** | | | | |
| **β_Intercept_** | 5.067 | 4.32 | 5.809 | **< 0.001** |
| **β_Age_** | 1.128 | 0.796 | 1.489 | **< 0.001** |
| **β_Female_** | 0.703 | 0.186 | 1.247 | **0.003** |
| $\boldsymbol{\sigma}$**_Cohort ID_** | 0.803 | 0.353 | 1.619 | — |

**Table S5.** **Posterior distribution of parameter estimates for potential predictors on the probability of not fledging (daily probability of not fledging from 16 days of age until 27 days).** The model included chick age (scaled to a mean of zero and standard deviation of 1), sex and their interaction as fixed effects, and Cohort ID and Mother ID as random effects. These parameters were intended to establish the effect these variables had on daily probabilities of not fledging for a given chick**.** Once the model for daily probability of not fledging was estimated from these effects, we derived the daily probability of fledging from the estimated trends as 1 – probability of a chick not fledging, for parameterizing the matrix model. The table provides the parameter estimates that are not back-transformed from the logit-link function. β_Intercept_ provides the estimated probability of not fledging for a male chick at an age of 21.5 days (i.e., the mean of the set 16 to 27 days). β_Age_ provides the estimated change in the probability of not fledging per standard deviation of age. β_Female_ provides the estimated difference in the probability of not fledging between female and male individuals at an age of 21.5 days. β_Age:Female_ provides the estimated interaction between chick age and sex. σ_Cohort ID_ and σ_Mother ID_ indicate the estimates for the between-cohort standard deviation and the between-mothers standard deviation, respectively. The table contains the mean, the 95% CrI (2.5% and 97.5% quantiles), the posterior probability of the hypothesis that the parameter β is smaller than zero. Probabilities in bold are comparable to a significant effect (P<0.05) according to frequentist statistics.

| **Parameter** | mean | 2.5% | 97.5% | P (β < 0) |
| --- | --- | --- | --- | --- |
| **β_Intercept_** | 0.195 | -0.712 | 1.06 | 0.292 |
| **β_Age_** | -3.099 | -3.744 | -2.521 | **> 0.999** |
| **β_Female_** | -0.612 | -1.181 | -0.055 | **0.985** |
| **β_Age:Female_** | 1.679 | 1.064 | 2.311 | **< 0.001** |
| $\boldsymbol{\sigma}$**_Cohort ID_** | 0.157 | 0.048 | 0.44 | — |
| $\boldsymbol{\sigma}$**_Mother ID_** | 0.053 | 0.003 | 0.128 | — |

**Table S6.** **Posterior distribution of parameter estimates for potential predictors on survival (daily chick survival probability within our free-living population from hatching until 11 days of age)** **of wild ruff chicks at Pitkänokka meadows, Liminka Bay, Finland.** The *Initial* *model* included chick age (scaled to a mean of zero and standard deviation of 1) as fixed effect and Cohort ID and Brood ID as random effects. These parameters were intended to establish the effect these variables had on wild survival**.** Because chick age did not have a clear effect, we removed it as an effect in the *Selected model* for parameterizing the matrix model. The table provides the parameter estimates that are not back-transformed from the logit-link function. β_Intercept_ provides the estimated wild survival of a male chick at an age of six days (i.e., the mean of the set 0 to 11 days). β_Age_ provides the estimated change in wild survival per standard deviation of age. σ_Cohort ID_ and σ_Brood ID_ indicate the estimates for the between-cohort standard deviation and the between-broods standard deviation, respectively. The residual standard deviation of the *Initial model* was 0.12 (0.08 – 0.27) and 0.09 (0.09 – 0.09) for the *Selected model*. The table contains the mean, the 95 credible interval (2.5% and 97.5% quantiles) and the posterior probability of the hypothesis that the parameter, β, is smaller than zero. Probabilities in bold are comparable to a significant effect (P<0.05) according to frequentist statistics.

| **Wild survival models** | | | | |
| --- | --- | --- | --- | --- |
| **Parameter** | mean | 2.5% | 97.5% | P (β < 0) |
| ***Initial model*** | | | | |
| **β_Intercept_** | 2.072 | -0.447 | 4.724 | **0.046** |
| **β_Age_** | -0.271 | -1.285 | 0.755 | 0.701 |
| $\boldsymbol{\sigma}$**_Cohort ID_** | 1.566 | 0.054 | 5.542 | — |
| $\boldsymbol{\sigma}$**_Brood_ _ID_** | 2.816 | 1.264 | 5.181 | — |
| ***Selected model*** | | | | |
| **β_Intercept_** | 2.23 | -0.034 | 4.637 | **0.026** |
| $\boldsymbol{\sigma}$**_Cohort ID_** | 1.471 | 0.04 | 5.316 | — |
| $\boldsymbol{\sigma}$**_Brood ID_** | 2.576 | 1.254 | 4.68 | — |

**Table S7. Posterior distribution of parameter estimates for tested predictors on fledging age**. Sex and adult body mass were modeled as fixed effects and Cohort ID and Mother ID as random effects. β_Intercept_ provides the estimated fledging age for males with a hypothetical adult body mass of zero grams. β_Female_ provides the estimated difference in fledging age between female and male individuals, and β_Adult mass_ provides the estimated change in fledging age per gram of adult body mass. σ_Cohort ID_ and σ_Mother ID_ indicate the estimates for the between-cohort standard deviation and the between-mothers standard deviation, respectively. Given are the mean, the 2.5%, the 97.5% quantiles, and the posterior probability of the hypothesis that the parameter is smaller than zero. The residual standard deviation was 1.46 (1.33-1.62). Probabilities in bold are comparable to a significant effect (P<0.05) according to frequentist statistics.

|  |  | Fledging age |  |  |
| --- | --- | --- | --- | --- |
| Parameter | mean [Days] | 2.5% [Days] | 97.5% [Days] | P (β<0) |
| β_Intercept_ | 26.97 | 23.48 | 30.50 | **< 0.001** |
| β_Female_ | -5.03 | -6.39 | -3.65 | **> 0.999** |
| β_Adult mass_ | -0.03 | -0.05 | -0.01 | **> 0.999** |
| σ_Cohort ID_ | 0.64 | -1.28 | 1.45 | — |
| σ_Mother ID_ | 0.44 | -0.86 | 0.89 | — |

**Table S8. Posterior distribution of parameter estimates for tested predictors on fledging age.** Sex and adult tarsus length were modeled as fixed effects and Cohort ID and Mother ID as random effects. β_Intercept_ provides the estimated fledging age for males with a hypothetical adult tarsus length of zero mm. β_Female_ provides the estimated difference in fledging age between female and male individuals, and β_Adult tarsus_ provides the estimated change in fledging age per mm of adult tarsus length. σ_Cohort ID_ and σ_Mother ID_ indicate the estimates for the between-cohort standard deviation and the between-mothers standard deviation, respectively. Given are the mean, the 2.5%, the 97.5% quantiles, and the posterior probability of the hypothesis that the parameter is smaller than zero. The residual standard deviation was 1.46 (1.32-1.61). Probabilities in bold are comparable to a significant effect (P<0.05) according to frequentist statistics.

|  |  | Fledging age |  |  |
| --- | --- | --- | --- | --- |
| Parameter | mean [Days] | 2.5% [Days] | 97.5% [Days] | P (β<0) |
| β_Intercept_ | 28.68 | 21.25 | 36.16 | **< 0.001** |
| β_Female_ | -4.18 | -5.44 | -2.90 | **> 0.999** |
| β_Adult tarsus_ | -0.13 | -0.27 | 0.01 | 0.965 |
| σ_Cohort ID_ | -0.72 | -1.38 | 1.53 | — |
| σ_Mother ID_ | 0.54 | -1.05 | 1.09 | — |

**Table S9. Posterior distribution of parameter estimates at sex-specific fledging ages.** Given are means [95% Cr].

| **Parameter** | **Males** | **Females** |
| --- | --- | --- |
| Wing length | 135 [133 – 136] | 112 [110 – 113] |
| Emerged primary length | 55.4 [53.9 – 57.1] | 45.6 [44.7 – 46.6] |
| Prop. of adult wing | 0.73 [0.72 – 0.74] | 0.73 [0.72 – 0.75] |
| Prop. emerged primary | 0.74 [0.72 – 0.75] | 0.73 [0.72 – 0.74] |
| Wing growth rate | 4.15 [3.74 – 4.55] | 3.98 [3.66 – 4.28] |
| Primary emergence rate | 4.72 [4.46 – 4.98] | 4.44 [4.18 – 4.71] |

**Table S10. Model response estimates for fledging sex ratios and selection coefficients from single-cohort matrix models.** The model response variables estimated in these two models were the post-fledge sex ratio (‘*Fledge* *SR*’), the change in SR from hatching to post-fledge ‘*ΔSR*’, and the sex-specific (♀: female; ♂: male) measures. These measures were the cross-sectional relative fitness *w* (calculated as post-fledge population proportion/hatching population proportion for each sex) and the selection coefficient *s* (calculated as the change in population/hatching population (1 – Fledging population proportion). The *Two-sex model* input parameters included the hatching sex ratio, sex-specific intrinsic survival, fledging advantage, and additive predation mortality. The *Unisex model* input parameters were the same as *Two-sex* model except intrinsic survival was the population average of both sexes. Ignoring sex-specific survival of chicks and hence relying on the *Unisex* model would have severely underestimated the sex ratio decline (*ΔSR*) and only accounted for 54% of the decline in comparison with the *Two-sex* model. Probabilities in bold are comparable to a significant effect (P<0.05) according to frequentist statistics. Table includes the response variables,$\theta$, their posterior mean, variance, 95% CrI (lower = 2.5 quantile; upper = 97.5 quantile) and the probability that the variable is lower than zero.

| **Response variable** | mean | variance | 2.5% | 97.5% | P ($\theta$ < 0) |
| --- | --- | --- | --- | --- | --- |
| ***Two sex model*** | | | | | |
| **Fledge SR** | 0.432 | 0.001 | 0.371 | 0.488 | **< 0.001** |
| ***ΔSR*** | -0.07 | <0.001 | -0.112 | -0.036 | **> 0.999** |
| $\boldsymbol{w}_{\boldsymbol{♀}}$ | 1.14 | 0.002 | 1.072 | 1.227 | **< 0.001** |
| $\boldsymbol{w}_{\boldsymbol{♂}}$ | 0.861 | 0.002 | 0.774 | 0.929 | **< 0.001** |
| $\boldsymbol{s}_{\boldsymbol{♀}}$ | 0.243 | 0.004 | 0.135 | 0.367 | **< 0.001** |
| $\boldsymbol{s}_{\boldsymbol{♂}}$ | -0.329 | 0.013 | -0.579 | -0.155 | **> 0.999** |
| ***Unisex model*** | | | | | |
| ***Fledge SR*** | 0.462 | <0.001 | 0.455 | 0.471 | **< 0.001** |
| ***ΔSR*** | -0.038 | <0.001 | -0.045 | -0.029 | **> 0.999** |
| $\boldsymbol{w}_{\boldsymbol{♀}}$ | 1.075 | <0.001 | 1.057 | 1.091 | **< 0.001** |
| $\boldsymbol{w}_{\boldsymbol{♂}}$ | 0.925 | <0.001 | 0.909 | 0.943 | **< 0.001** |
| $\boldsymbol{s}_{\boldsymbol{♀}}$ | 0.14 | <0.001 | 0.109 | 0.166 | **< 0.001** |
| $\boldsymbol{s}_{\boldsymbol{♂}}$ | -0.163 | <0.001 | -0.199 | -0.122 | **> 0.999** |

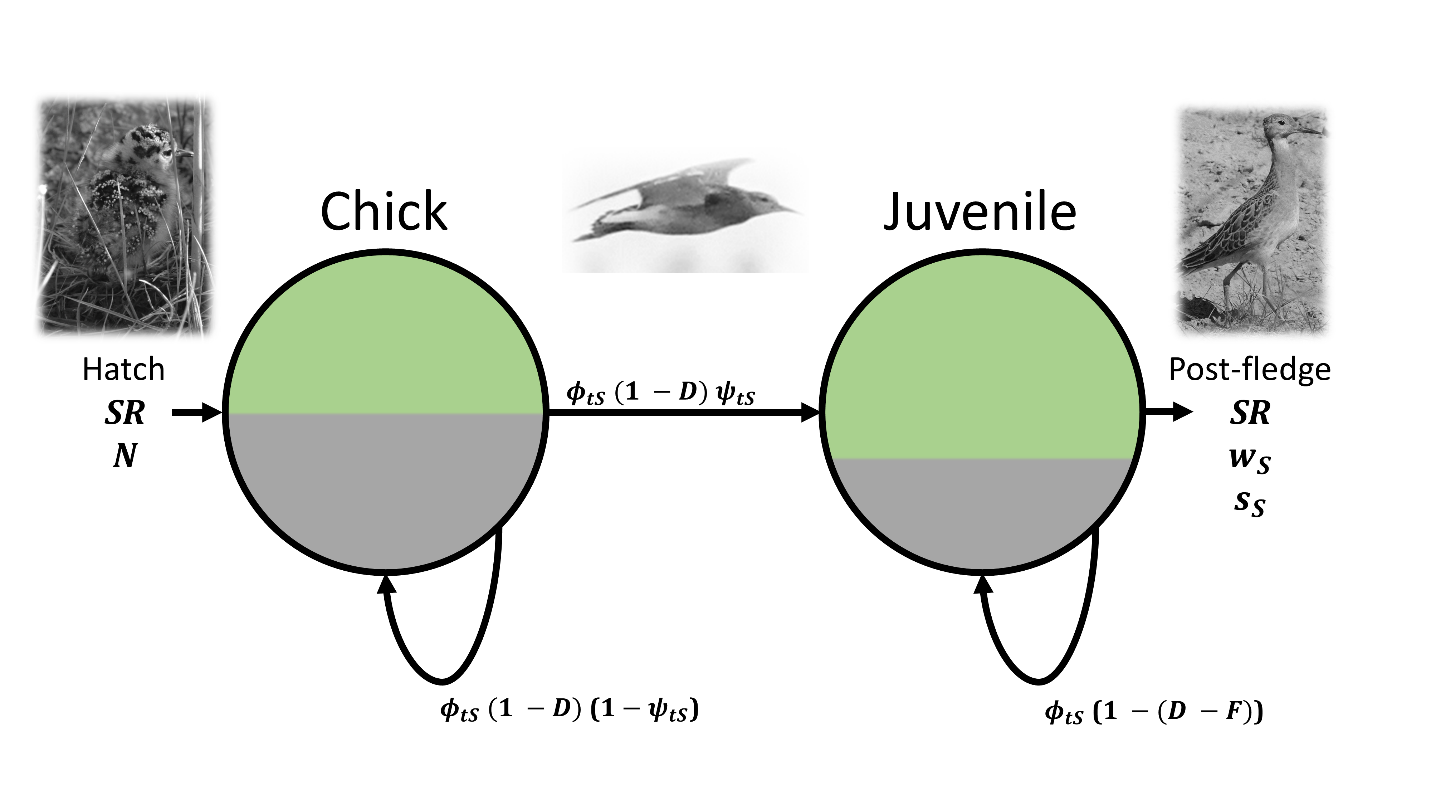

**Fig S1. Flow diagram representing single-cohort matrix model to estimate sex ratio (SR) change from hatching to post-fledging (age 27 days).** Color shades within life-stages (chick and juvenile) denote the hypothetical relative proportions of males (gray) and females (green). The diagram illustrates the processes of intrinsic survival (*ϕ*) and fledging probabilities (*ψ*) between the life-stages, when additive predation mortality (*D*) occurs and juveniles escape predation with a fledging advantage (*F*). *ϕ* and *ψ* are sex- and age-specific variables denoted with the indices, sex (*S*) and time (*t*) respectively. *N* chicks hatch in each cohort and the hatching sex ratio (HSR) determines the proportion of males (*HSR*) and females (1 – *HSR*). After hatch, chicks survive at a rate of $\phi_{ts}(1-D$) and fledge to become juveniles at a rate of $\psi_{s}$. Once fledged, juveniles survive at a rate of $\phi_{ts}(1-(D-F$)) until all chicks in the cohort have become juveniles. At this point the cohort reaches the post-fledge life stage and the post-fledge sex ratio (post-fledge SR), sex-specific cross-sectional relative fitness (*w*) and selection coefficients (*s*) can be calculated.

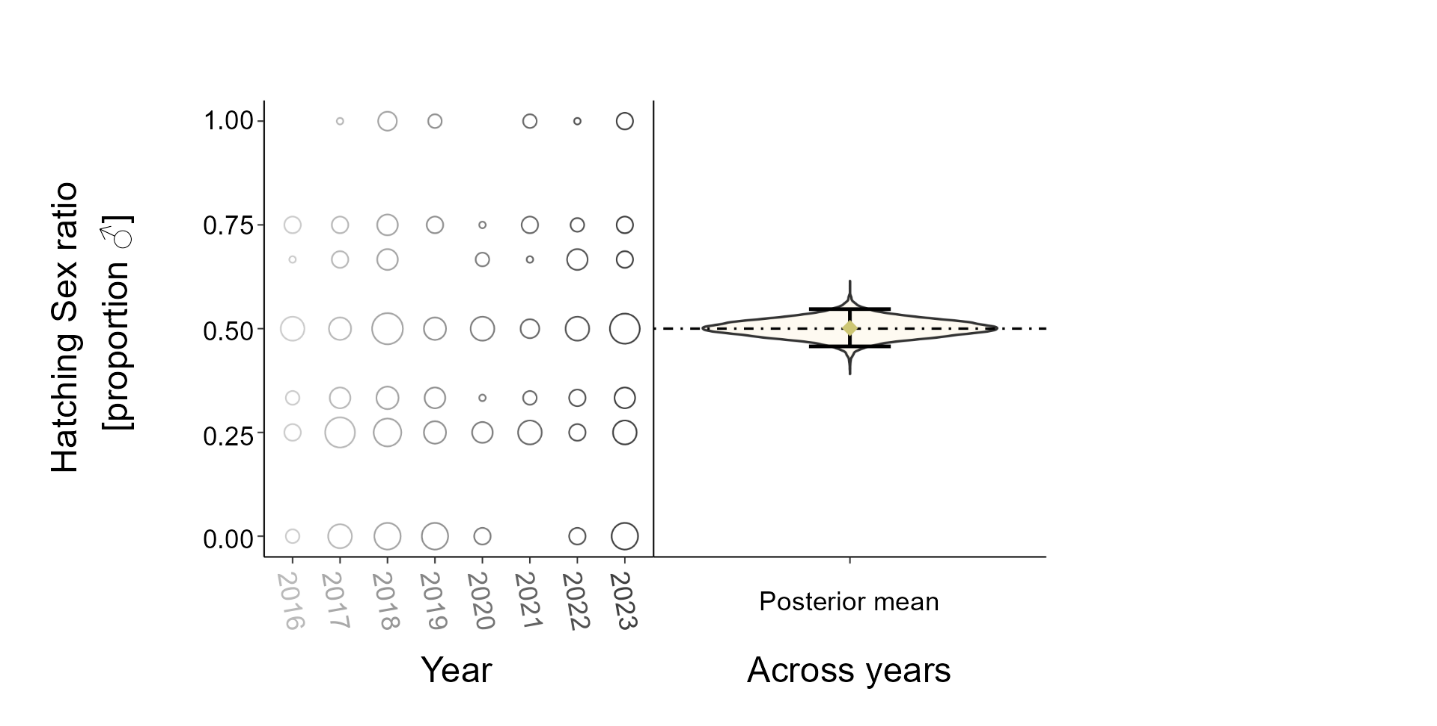

**Fig S2. Hatching sex ratios (proportion of male chicks) of wild Ruff clutches collected at Pitkänokka meadows, Liminka Bay, Finland from 2016 until 2023.** Left, sex-ratios of sampled chicks with circle size representing the relative number of clutches/broods within a year that have a given sex ratio. Right, the posterior density, violin plot of a Bayesian Generalized Mixed Effects Model where the response was the probability of a chick sampled being male. Besides an intercept (mean sex ratio per year, per clutch) there were no fixed effects in this model, however the model included Cohort ID, the year in which the sample was taken, and Clutch ID, the identity of the clutch the sample belonged to. Broken horizontal line is set at 0.5, the golden diamond is the posterior mean across clutches and years, and the black vertical lines are the 95% credible intervals. The posterior standard deviation was 0.32 (0.31 – 0.32).

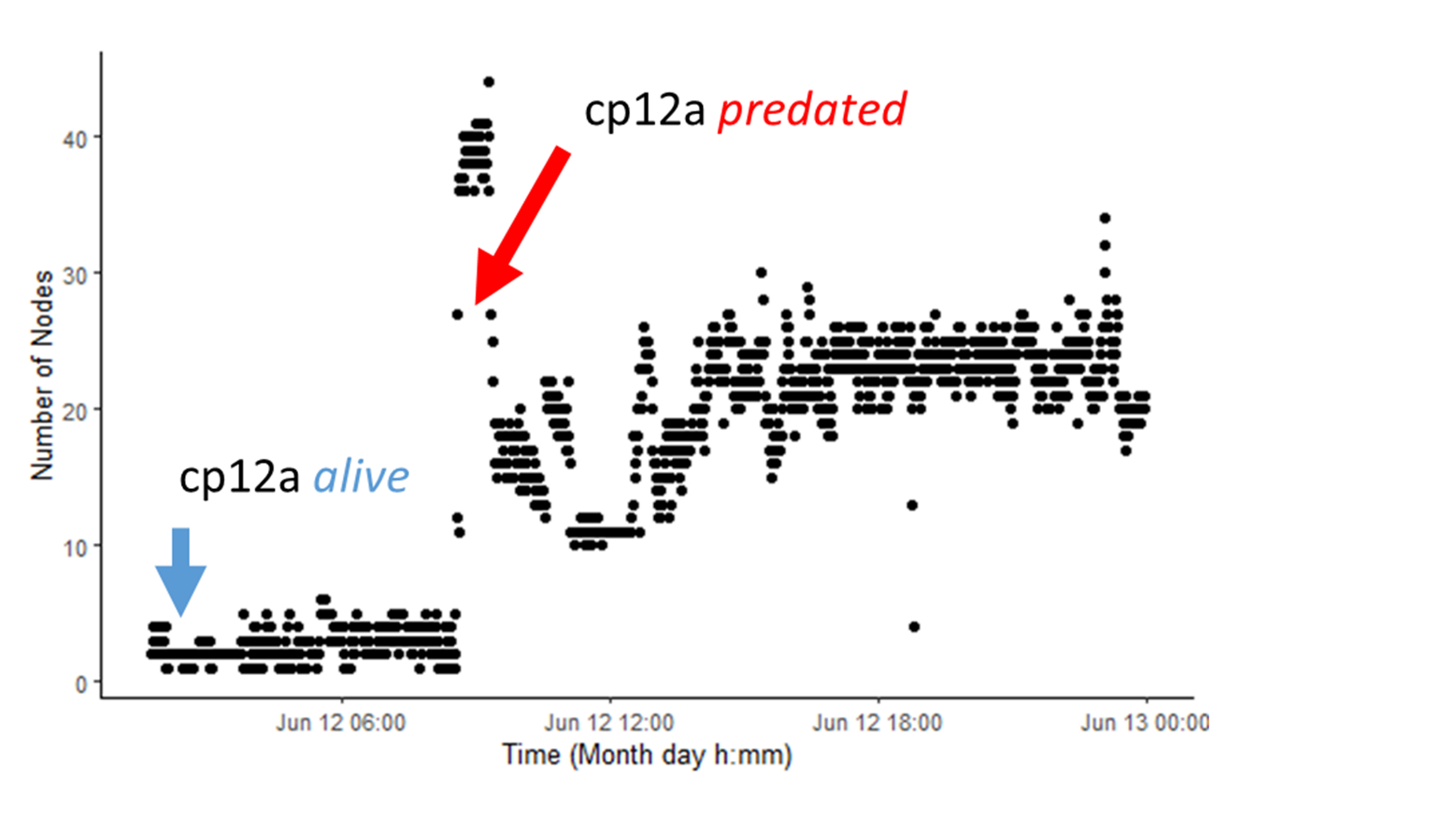

**Fig S3. Example of remote tracking data of radio tagged wild Ruff chicks** **at Pitkänokka meadows, Liminka Bay, Finland.** An example of how we determined chick fate. The chick cp12a can be seen alive with relatively few nodes detecting the chick. However, there is an abrupt change in the number of nodes detecting the chick’s radio tag indicating that the chick was predated. It was predated by a gull who swallowed the tag. The gull was flying with the ingested tag frequently leading to detection of many more nodes than whilst the chick was in the vegetation when it was alive (before 12 June 9 am). In addition to these data, we confirmed the chick’s predation with a hand radio receiver.

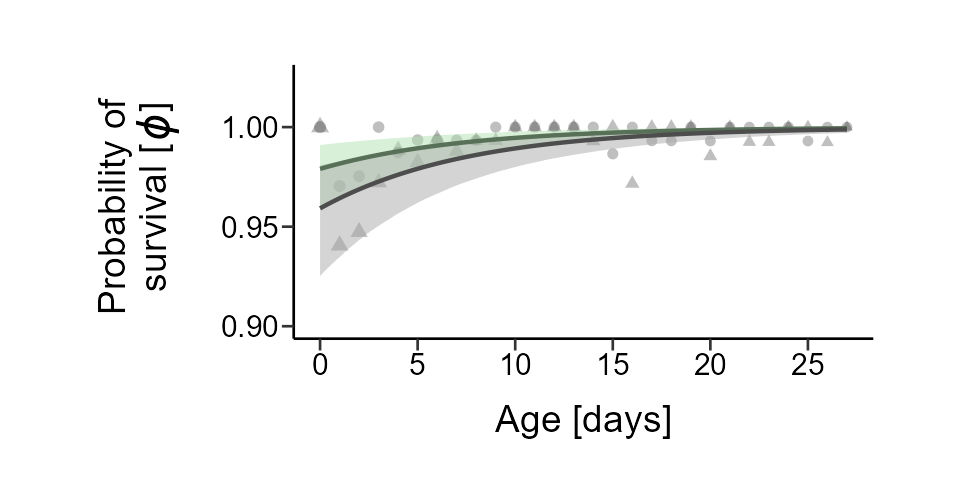

**Fig S4. Sex-specific intrinsic survival probabilities changes with age in captive Ruff chicks. Survival was model with data from hand raised Ruff chicks from 2018 until 2023.** Male survival shown in gray, female survival in green. Lines are the posterior mean probability of survival and shaded areas are the 95% credible interval (*Φ*), from a Bayesian Mixed Effects Daily Survival model. Although age was standardized for fitting the model and estimating the posterior response, here we display the non-standardize age in days to ease interpretation, however the *y*-intercept in the model was set at the mean age in the set of 0 to 27 (13.5 days old). Points are the proportional outcomes of observed Ruff chicks that survived divided by the total number of chicks observed at that age. Points are sex-specific proportions of observed chicks that survived to the next day (triangles = males; circles = females) and size of the point represents the relative number of chicks observed at that age. The model included the random effect Cohort ID, the year in which the observed chick hatched. Although age was standardized for the model, here we display the non-standardize age in days to ease interpretation. The residual standard deviation was 0.01 (0.01 – 0.02).

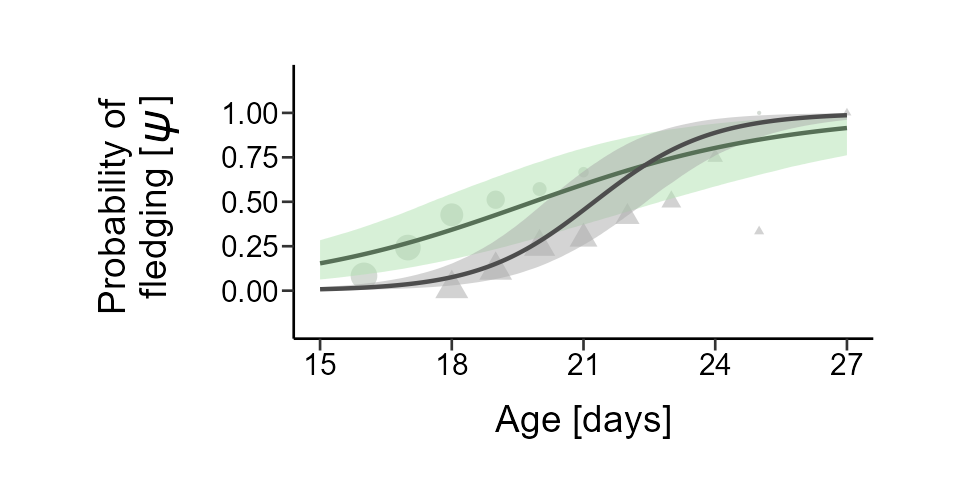

**Fig S5. Sex-specific probability of fledging at a given age in captive Ruff chicks. Fledging age was modeled with data from fledge tests using hand raised Ruff chicks from 2018 until 2023.** Male fledging probability shown in gray, female fledging probability in green. Lines are the posterior mean and shaded areas are the 95% credible interval of the probability of fledging (*ψ*) from a Bayesian Mixed Effects Daily Survival, where 'not fledging' was analogous to 'survival' and *ψ* was derived from the estimated response as '1 – the probability of not fledging'. Although age was standardized for fitting the model and estimating the posterior response, here we display the non-standardized age in days to ease interpretation, however the *y-*intercept in the model was set at the mean age in the set of 16 to 27 (21.5 days old). Points are the proportional experimental outcomes of tests conducted on Ruff chicks that flew divided by the total number of chicks tested at that age. Points are sex-specific proportions (triangles = males; circles = females) and size of the point represents the relative number of chicks tested at that age. The model contained the random effects Cohort ID, year in which the tested chick hatched, and Mother ID, the mother ID of the tested chick. The residual standard deviation was 0.31 (0.27 – 0.34).

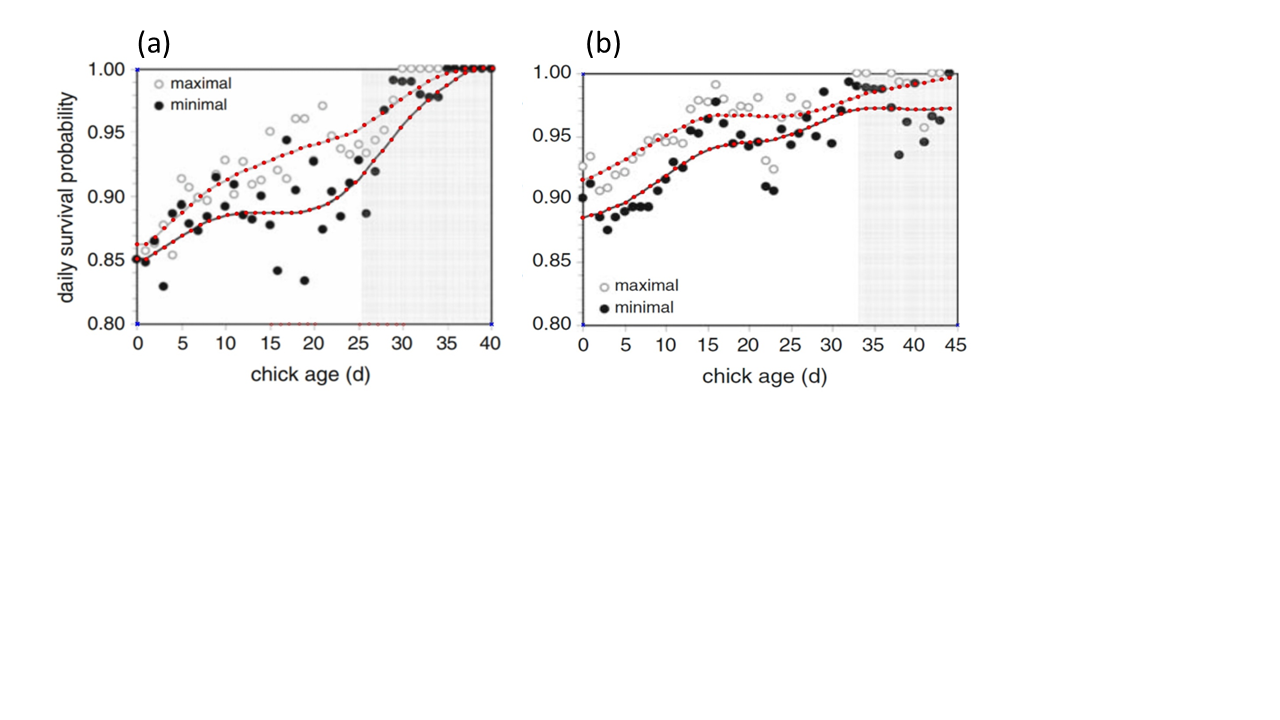

**Fig S6. Historic survival data of shorebird chicks used to calculate the fledging advantage for Ruff juveniles.** Survival data were extracted from Schekkerman et al. (2009) Fig. 2. (a) Black-tailed Godwit, *Limosa limosa*, chick survival. (b) Northern Lapwing, *Vanellus vanellus*, chick survival. Black dots are the observed proportion of chicks with known fates and unknown fates. Here unknown fates were recorded as mortalities. Open circles by contrast, are the observed proportion of chicks surviving with only known fates. Black and white trend lines are estimated daily survival from black dot data and open circle data respectively. Gray shaded areas start at the fledging date for the species. Red dots are points we extracted using the digitize package (Poisot 2011) in R. Red dots before the gray shaded area correspond to pre-fledging survival estimates whereas red dots in gray shaded areas refer to post-fledging survival estimates.

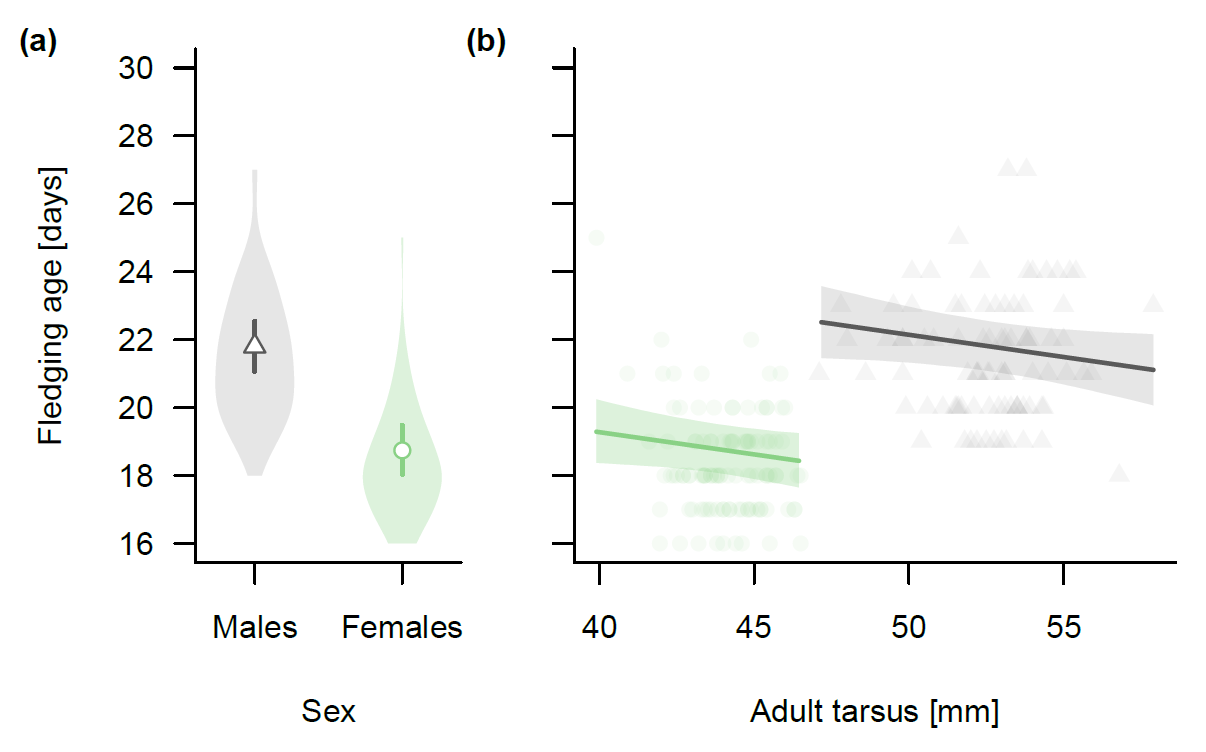

**Fig S7.** **Variation in fledging age by sex and adult tarsus length. (a)** Fledging ages of male and female chicks when statistically controlled for cohort and mother ID. The probability of females (green) fledging earlier than males (gray) was higher than 0.99. Means (triangle for males and circle for females) and 95% credible intervals (lines) correspond to fledging ages at a sex-specific mean tarsus length (44.1mm in females and 52.6mm in males). Shaded areas reflect the density distribution of raw data. **(b)** Fledging ages according to final tarsus length as adults for male and female chicks. Given are the mean and 95% credible intervals fledging ages (line and shaded area). The probability of individuals with longer tarsi fledging earlier than individuals with shorter tarsi within sex was 0.965.

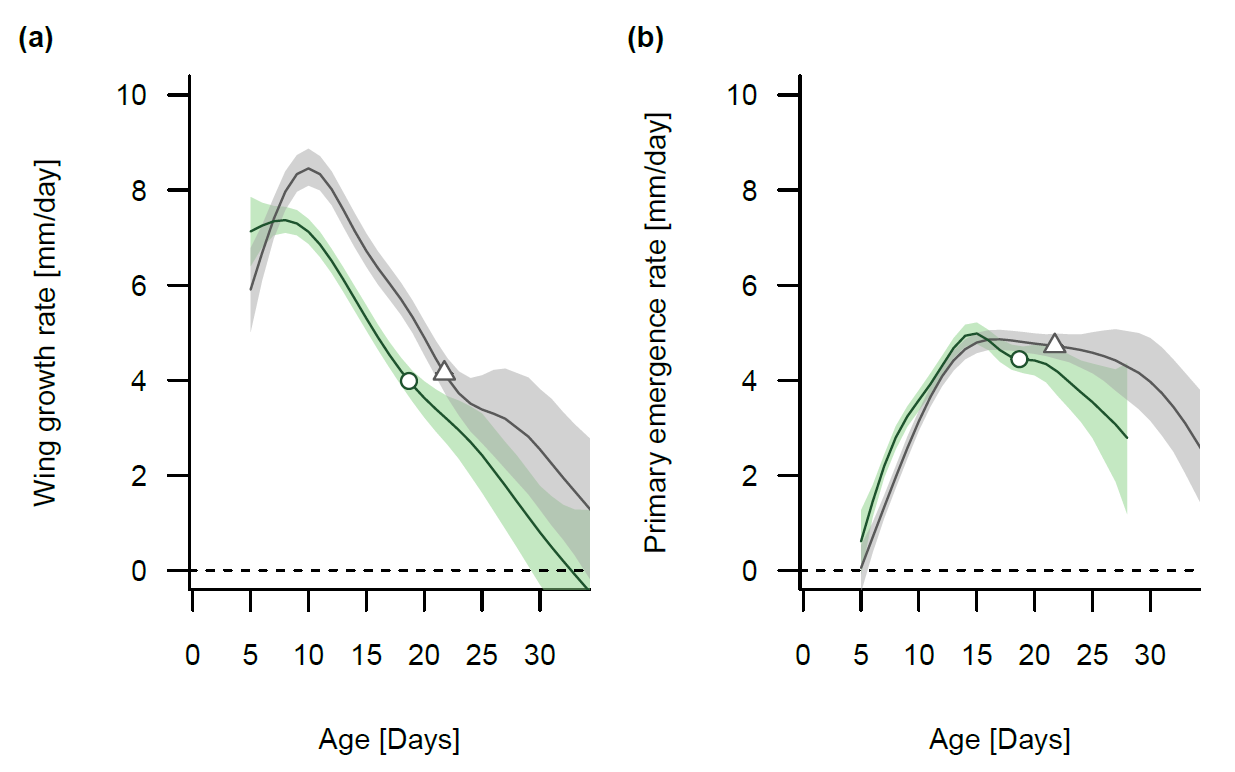

**Fig S8. Wing and primary growth rates in male and female Ruff chicks that were hand raised in captivity.** (a) Wing growth rate at a given age. (b) Primary emergence rate at a given age. Means and 95% credible intervals (lines and shaded areas) of males are in gray and of females in green. Open symbols indicate wing growth rate (a) and primary emergence rate (b) at fledging for males (triangles) and females (circles).

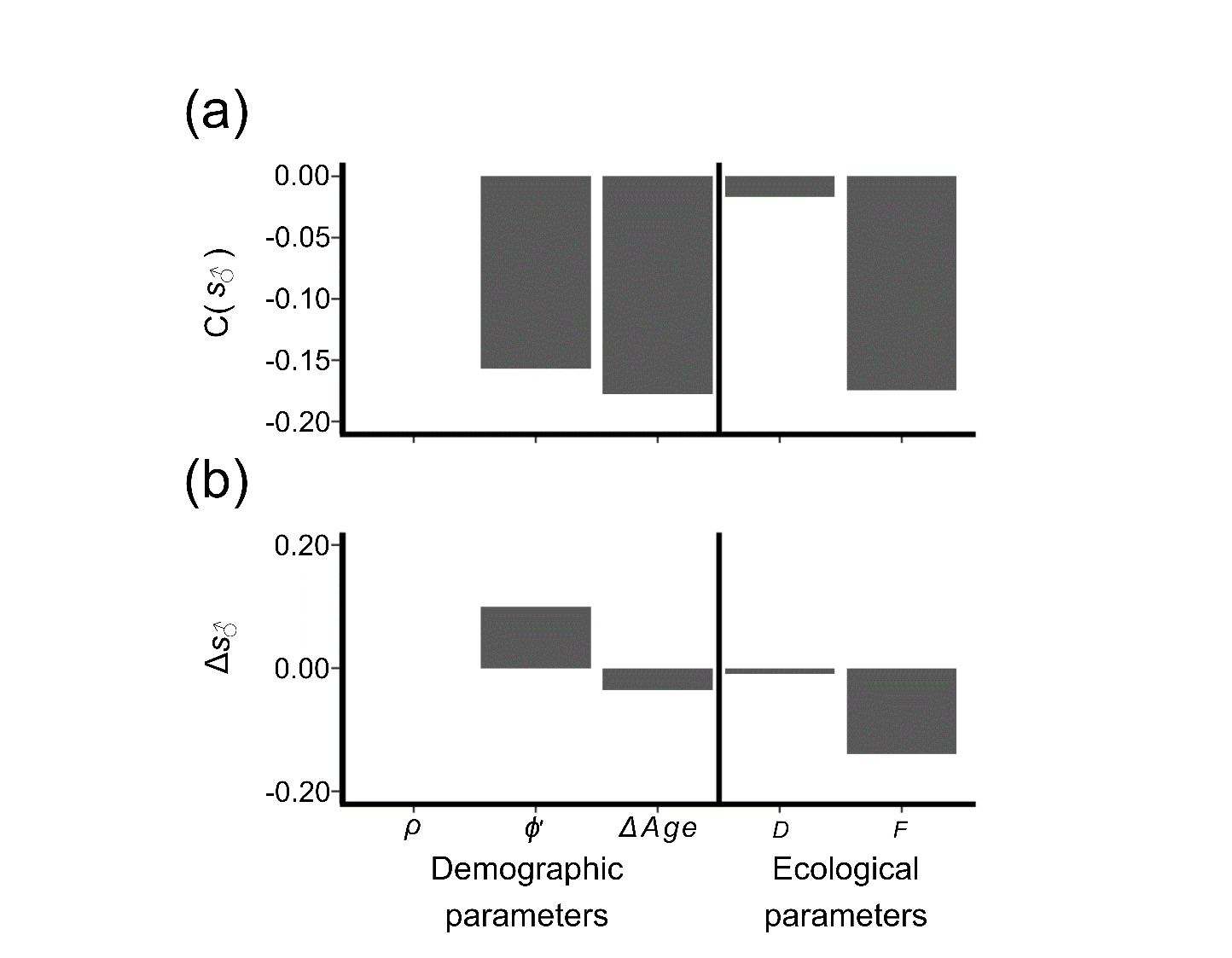

**Fig. S9. Sensitivity analyses showing the contribution of mean parameters estimates to projected male selection coefficient(**$\boldsymbol{s}_{\boldsymbol{♂}}$**) at the mean parameter estimate (a), or by an increase of one standard deviation (d).** Left, demographic parameters: hatching sex ratio (ρ), cumulative intrinsic survival (*ϕ'*), and sex difference in fledging age (*ΔAge*). Right, ecological parameters: additive predation mortality (*D*), and fledging advantage (*F*).

### References

1.

Dushoff, J., Kain, M.P. & Bolker, B.M. (2019). I can see clearly now: reinterpreting statistical significance. *Methods in Ecology and Evolution*, 10, 756-759.

2.

Eberhart-Phillips, L.J., Küpper, C., Miller, T.E.X., Cruz-López, M., Maher, K.H., dos Remedios, N. *et al.* (2017). Sex-specific early survival drives adult sex ratio bias in snowy plovers and impacts mating system and population growth. *Proceedings of the National Academy of Sciences*, 114, E5474.

3.

Evans, P.R. (1986). Correct measurement of the wing-length of waders. *Wader Study Group Bulletin*, 48, 11.

4.

Giraldo-Deck, L.M., Goymann, W., Safari, I., Dawson, D.A., Stocks, M., Burke, T. *et al.* (2020). Development of intraspecific size variation in black coucals, white-browed coucals and ruffs from hatching to fledging. *Journal of Avian Biology*, 51.

5.

Giraldo-Deck, L.M., Loveland, J., Goymann, W., Tschirren, B., Burke, T., Kempenaers, B. *et al.* (2022). Intralocus conflicts associated with a supergene. *Nature Communications*, 13, 1384.

6.

Grant, A.S. & Oring, L.W. (1997). *Guidlines to the use of birds in research* Third Edition 2010 edn. Ornithological Council, Washington, D. C.

7.

Korner-Nievergelt, F., Roth, T., von Felten, S., Guélat, J., Almasi, B. & Korner-Nievergelt, P. (2024). Daily nest survival. In: *Bayesian data analysis in ecology using R and Stan* bookdown.

8.

Lees, D., Schmidt, T., Sherman, C.D.H., Maguire, G.S., Dann, P., Ehmke, G. *et al.* (2019). An assessment of radio telemetry for monitoring shorebird chick survival and causes of mortality. *Wildlife Research*, 46, 622-627.

9.

Mason, L.R., Smart, J. & Drewitt, A.L. (2017). Tracking day and night provides insights into the relative importance of different wader chick predators. *Ibis*, 160, 71-88.

10.

Schekkerman, H., Teunissen, W. & Oosterveld, E. (2009). Mortality of black-tailed godwit *Limosa limosa* and Northern lapwing *Vanellus vanellus* chicks in wet grasslands: influence of predation and agriculture. *Journal of Ornithology*, 150, 133-145.

11.

Umlauf, N., Klein, N. & Zeileis, A. (2018). BAMLSS: Bayesian additive models for location, scale, and shape (and beyond). *Journal of Computational and Graphical Statistics*, 27, 612-627.
